# Trophic Interactions of *Anopheles Gambiae* Mosquito Larvae in Aquatic Ecosystem: A Metagenomics Approach

**DOI:** 10.1101/2024.11.01.621049

**Authors:** Afia Serwaa Karikari, Jewelna Akorli, Francis Gbogbo, Ignatious Cheng Ndong, Akosua Bonsu Karikari, Akua Afriyie Karikari, Rosina Kyerematen, Akwasi Agyenkwa Mawuli, Thomas Gyimah, Fred Aboagye-Antwi

## Abstract

**Background:** Understanding the trophic interactions of *Anopheles gambiae* mosquito larvae in aquatic environments is essential for malaria prevention efforts such as in gene drive application. In this study, metagenomics method was employed to explore the feeding behavior and ecological associations of *An. gambiae* larvae within their aquatic ecosystems.

**Methods:** *Anopheles gambiae* larvae and their co-existing fauna species were collected from December 2022 to June 2023. DNA from dissected midguts of the co-existing fauna was sequenced using Illumina NovaSeq for diet analysis through a shotgun sequencing approach.

**Results:** The study revealed a complex network of trophic interactions in the freshwater habitat, with significant resource sharing between *A. gambiae* larvae and other filter-feeding species, including various Diptera and non-Dipteran feeders. Contrary to assumptions, predator species did not exhibit exclusive predation on *Anopheles* larvae, preferring other fauna instead.

**Conclusion:** This study has demonstrated complex trophic interactions among *Anopheles gambiae* larvae and other organisms in freshwater ecosystems. It has offered essential insights for optimizing vector control strategies, such as in gene drive applications. Additionally, the study has provided valuable information on the aquatic fauna of the study area, which can serve as a baseline for developing a macroinvertebrate identification database for freshwater systems in Ghana. A complementary study that further explores the ecological role of *An. gambiae* larvae in these freshwater habitats is presented in a subsequent publication.

## Background

Malaria, a severe infectious disease, continues to impose a significant global health burden, with the *Anopheles gambiae* mosquito serving as a primary vector [1-3]. Recent data indicates that approximately 608,000 people died out of 247 million reported global malaria cases [4]. Ghana accounts for 2.2% of worldwide malaria infections and 2% of related deaths, placing it among the top 15 most affected nations. Furthermore, it is responsible for 4% of the malaria burden in West Africa [4]. To mitigate this burden, the country implemented the “high burden, high impact” strategy in November 2019, and used mass campaigns to apply indoor residual spraying (IRS) and insecticide-treated nets (ITNs) [5]. However, behavioral issues and low acceptance have compromised the efficiency of these interventions. For instance, many residents report difficulties hanging rectangular nets and prefer alternative methods such as mosquito sprays [5].

Along with artemisinin-based combination therapy, ITNs and IRS are widely used as primary malaria control techniques worldwide and are not exclusive to Ghana [3]. Nevertheless, global malaria eradication efforts are facing setbacks due to rising resistance in both parasites and mosquito vectors against drugs and insecticides [4, 6]. Compounding these challenges is the lack of an effective large-scale malaria vaccine and the tendency of mosquitoes to bite outdoors, both of which sustain malaria transmission [7]. There is a need to develop safer, more efficient, cost-effective and eco-friendly vector control than insecticides [8].

Recent progress in genetic engineering has led to the creation of novel techniques for genome modification of various species to offer safer, cost-effective, and environmentally sustainable alternatives to insecticides [8]. Among these innovations, gene drive technology shows promise by allowing the use of genetically modified mosquitoes aimed at controlling malaria and other vector-borne diseases [9]. One prominent gene drive strategy under consideration involves population suppression, which seeks to reduce the population of malaria-transmitting mosquitoes over time [10].

Despite its potential, gene-drive technology has generated substantial debate over biosafety and ecological risks [7,10, 11, 12]. Concerns revolve around unintended ecological disruptions, such as competitive advantage, secondary extinctions, and *Anopheles gambiae’s* ecological significance within ecosystems [10, 11]. To evaluate these potential risks requires an in-depth knowledge into the mosquito species and the relationship with their community [12].

In recent times, there are proposals to focus on immature mosquitoes, either independently or as part of a multifaceted vector control strategy, to reduce or eliminate malaria [13-16]. Unlike the adult mosquitoes, which are highly mobile and thrives in varied environments (including soil, rocks, and vegetation), the larvae are confined to aquatic habitats, making them a more manageable stage [17]. Targeting the larvae could disrupt the mosquito’s development earlier in the life cycle, complementing adult control measures within integrated vector management systems [13-16]. Achieving success at larval control requires a detailed understanding of their ecology, particularly the trophic interactions, such as food sources, competitors, and predators within their aquatic habitats. Unfortunately, the ecology of malaria vectors, especially the feeding interactions remains poorly understood [18].

Food acquisition is a critical activity for all living organisms since it affects the energy available for maintenance, growth, and reproduction [19,20]. It also influences species interactions and the structure and function of food webs [19]. Relating food to trophic network interactions requires tools to examine a species’ dietary composition. Gut content analysis has been a common and useful technique for investigating animal diet components and feeding ecology [19]. Contemporary studies employed gut content analyses and polymerase chain reaction (PCR) methods, to identify arthropod prey within predator guts [21,22]. Nonetheless, the versatility and application of PCR-based methods have several limitations. One such challenge is that the commonly used barcode markers, like cox1 and rRNAs, lack sufficient conservation across different taxonomic groups, compounding the creation of universal or broad-range primers without target-matching limitations. Consequently, marker selection and primer efficiency can introduce biases, potentially skewing species identification and leading to inaccurate estimates. of the abundance of certain organisms [26 -30]. Recent advances in metagenomics offer a more precise and comprehensive approach. The technique provides greater taxonomic resolution, allowing for the detection of prey species, endoparasites, and associated microbiomes by matching unassembled DNA reads from samples against reference databases [31,32]. Therefore, the accuracy of these analyses, depends on the availability and quality of DNA libraries [32].

Ghana currently lacks a comprehensive DNA reference library for its aquatic macroinvertebrates, hence external datasets with well-established repositories [33-34] were employed in this study, to explore the trophic interactions of *An. gambiae* larvae through metagenomics approach. Specifically, the study aimed to identify the food resources and probable competitors and predators of *Anopheles gambiae* larvae in their natural habitats. The findings will contribute to knowledge essential for developing innovative mosquito control strategies, including gene drives. In addition, by producing a classification of aquatic macroinvertebrates through gut composition analysis, the results of the study would provide a first-hand insight into the aquatic macroinvertebrates of Ghana. This would be informative in developing an eDNA protocol for monitoring probable freshwater macroinvertebrates in the country.

The second phase of this research which involves ecological network analysis to investigate the broader ecological roles of *An. gambiae* larvae will be presented in a subsequent publication.

## Materials and Methods

### Study Areas

This study was conducted in two rural communities: Mafi Agorve and Abutia Amegame, both situated in the Volta Region of Ghana (Fig. 1). The Volta Region lies approximately 2.5 hours by road from Accra, the capital city of Ghana, and covers a total land area of 20,572 km^2^, located between 3°45′ and 8°45′ N latitude. The study sites comprise of savannah and woodland habitats, where two primary members of the *Anopheles gambiae* complex are found: *Anopheles gambiae* sensu stricto; commonly associated with forested areas and *Anopheles coluzzii*, which is more frequent in savannah landscapes and areas impacted by human activity [35].

**Fig 1.**
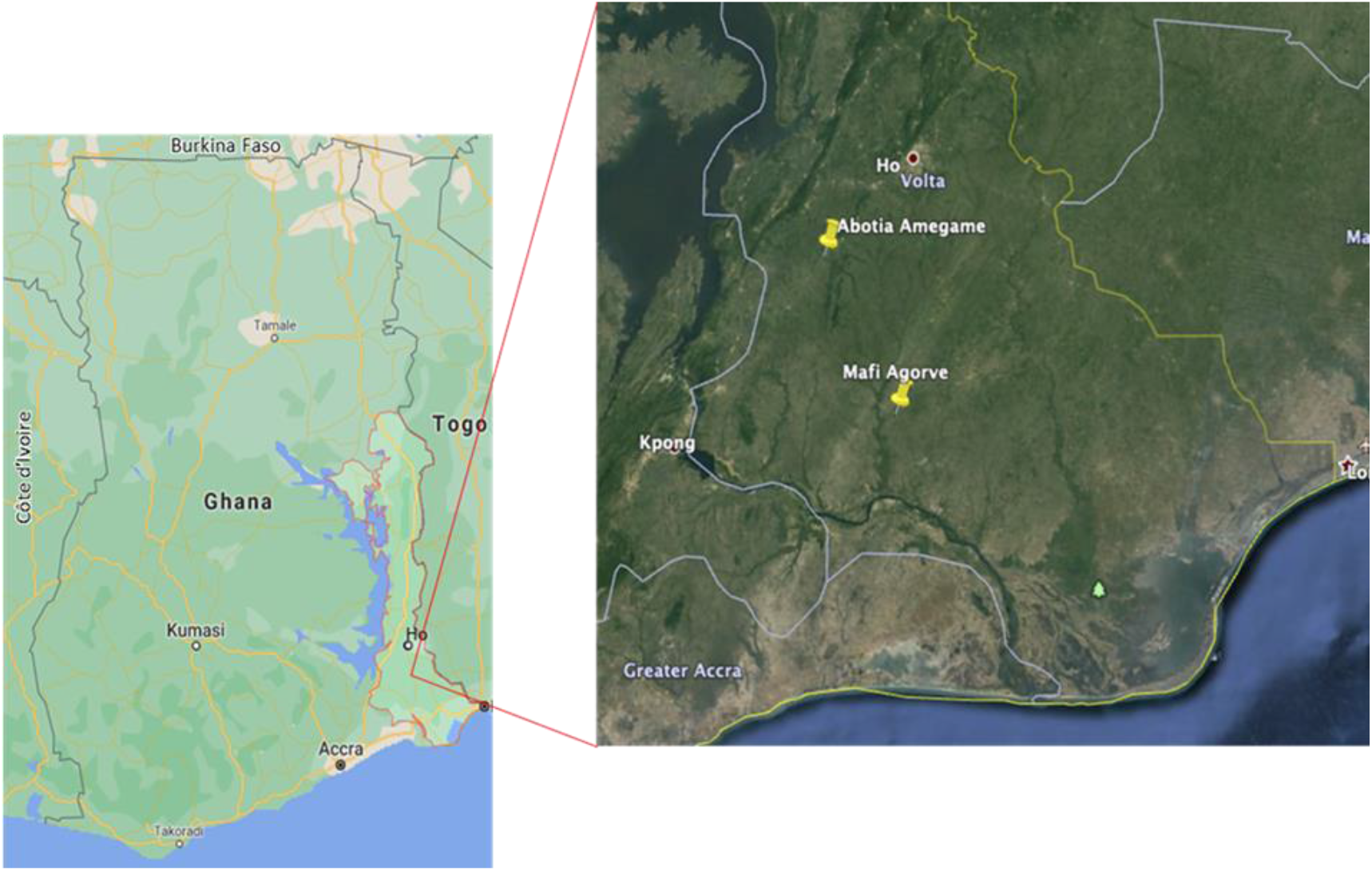
Map of Ghana showing the study sites in the Volta Region

Both areas experience a bimodal rainfall pattern, with a major rainy season from mid-April to July and a minor season from September to November. However, rainfall varies significantly between the two sites. Mafi Agorve, belonging to the Central Tongu district of the region, receives an average annual rainfall of 1,100 mm, though it remains insufficient for optimal crop production [36]. In contrast, Abutia Amegame which is part of the Ho municipality, located farther north is much drier, with an average of 192 mm of rainfall annually [36].

Temperatures across both areas remain relatively stable throughout the year. In Mafi Agorve, the minimum temperature is around 22°C, with March being the hottest month and July and August the coolest. The region also maintains high relative humidity, averaging 80%, creating favorable conditions for both farming and mosquito breeding [36, 37]. Abutia Amegame experiences slightly higher temperatures, ranging between 22°C and 32°C [36].

The combination of savannah and woodland ecosystems across both locations provided an ideal setting for examining the larval ecology of *An. gambiae* under varying environmental conditions.

### Fauna species used for the study

The study examined nine faunal species from five orders: Diptera, Hemiptera, Calanoida, Cyprinodontiformes, and Odonata. The selected species included *Anopheles* mosquito larvae and other dipterans such as *Culex, Clunio*, and *Toxorhynchites*, the latter also serving as a predator in this study. Non-dipteran species involved in the analysis included Notonecta (Hemiptera), *Ischnura* and *Pantala* (Odonata), *Gambusia* fish (Cyprinodontiformes), and a copepod (Calanoida). These non-dipteran species were selected based on documented evidence of their predatory behavior [38–42] and ecological associations with *Anopheles gambiae* larvae in the study areas. Upon collection, each specimen was preserved in separate vials containing 95% ethanol and stored at -80°C until gut content dissection.

### Gut content dissection

Dissections and DNA extractions were conducted at the Department of Parasitology, Noguchi Memorial Institute for Medical Research at the University of Ghana. The methodology followed stringent protocols to ensure accuracy and prevent contamination. All surfaces were thoroughly cleaned with a 70% alcohol solution. Dissecting pins and Petri dishes were serially sterilized using 70% alcohol, 10% bleach, and 1% phosphate-buffered saline (PBS) before each dissection session to maintain aseptic conditions.

The dissections were performed using a Motic K Series dissecting microscope with a magnification range of 6x - 12x, guaranteeing detailed observation. Each specimen also underwent a triple wash with PBS before each dissection, to eliminate any external contaminants. For insect species, precise excision of the head, and appendages, clipping of the 5th-7th abdominal segments, and removal of the remaining abdominal covering were made to expose the midgut. The midgut, containing the gut contents, was then carefully transferred to a sterilized slide and washed with PBS before being pipetted into a 2 ml Eppendorf tube also containing PBS. For fish species, the gut was meticulously removed from the slit near the operculum. Conversely, copepods were utilized entirely for DNA extraction due to their relatively smaller size. The samples were stored at -80°C until DNA extraction to preservation.

### DNA extraction

During the DNA extraction process, excess PBS was meticulously pipetted from the samples. Subsequently, an average of 10 gut samples were combined per species per site, except for *Gambusia* sp and the copepod, for which only one specimen was available due to insufficient samples. The combined samples were homogenized in the ZYmoBiomics DNA Kit (lysis buffer) using a plastic pestle attached to a hand-held homogenizer. Following homogenization, DNA was extracted using the ZYmoBiomics DNA Kit, following the manufacturer’s standard proteinase K protocol. The extraction buffer was employed for mock dissection extraction to serve as a negative control in the process

### Library Construction, Quality Control and Sequencing

Before sequencing, the genomic DNA (gDNA) samples were taken through two primary quality control (QC) methods: Nanodrop was used for preliminary quantitation of the DNA subsequent to an Agarose Gel Electrophoresis to assess DNA degradation and identify potential contamination.

Following successful quality control (QC) checks, the DNA in the samples was enriched with oligo (dT) beads after which the enriched gDNA was randomly broken into short fragments of approximately 350 bp by the addition of a fragmentation buffer. The resulting fragments underwent end-repair, A-tailing, and ligation with Illumina adapters. The adapter-attached fragments or libraries were subjected to size selection, PCR amplification, and purification prior to another three quality control checks involving Qubit 2.0 Pre library concentration testing, an Agilent 2100 insert size assessment and a qPCR precise quantification of the library’s effective concentration.

The quantified libraries were combined and introduced into Illumina sequencers based on the effective library concentration and the required data amount. All gut samples underwent sequencing on the Illumina NovaSeq platform, generating 150 bp paired-end reads at a depth of 6Gb per sample. The reads were assessed through a quality control check with a minimum Phred quality score of 20. Retained reads after quality control were then converted to Fasta format.

### Data Analyses

#### Reference Database

A seqTaxDb taxonomic database known as: “METSP_zenodo_3247846_uniclust90_2018_08_seed_valid_taxids.tar.gz” [43] was used for the study. This database has approximately 88 million entries and comprises of two main resources: Uniclust90 seed proteins which is a database of clustered and deeply annotated protein sequences that groups UniProtKB sequences based on their pairwise sequence identity at the level of 90% [44] and proteins from the *Marine Microbial and Eukaryote Transcriptome Sequencing Project (MMETSP)* which combine 678 microbial eukaryotic reference transcriptomes [45, 46]. The combined database was downloaded from the Zenodo repository for use. In addition, a Kraken2 run was done to find if the unmapped reads from the eukaryotic profiling were prokaryotes.

#### De novo assembly

Quality scores across the Illumina paired-end reads were examined using fastqc [47] and MultiQC [48], and poor-quality reads were trimmed using fastp [49]. After this quality control, reads from each sample were assembled into contigs using the metagenomic assembler, Megahit [50] under default settings with k-max = 141. The results were a FASTA file for each sample containing metagenomic contigs.

#### Taxonomic assignment to metagenomic contigs

MMseqs2 taxonomy mode was combined with the database to classify the contigs in each FASTA file. MMseqs2 is a novel tool for tagging metagenomic contigs [51]. It extracts all possible protein fragments from each contig, quickly retains those that can contribute to taxonomic annotation, and assigns them robust labels using Bayesian-based Least Common Ancestor [52], known as 2bLCA, which determines the contig’s taxonomic identity by weighted voting [53].

Assignment occurred following Mirdita *et al*, as follows: Each query fragment q was searched against the reference database, producing a list l of homologous target sequences. The aligned region between q and its top hit t with an E-value E(q, t) was then aligned against all other targets in l. The fragment q was assigned the Lowest Common Ancestor (LCA) of the taxonomic labels for all targets with an E-value below E(q, t). To optimize efficiency, the list l was realigned to avoid a second search during the 2bLCA process. Each query q contributed a weight of -log E(q, t) to its corresponding taxonomic label and all higher-level labels up to the root. The final taxonomic assignment for a contig was determined by the most specific label that satisfied the majority rule. The support for each label was calculated as the sum of contributing weights divided by the total weight across all assigned labels.

In this run, the LCA-mode was left at the default value of 3 but the report-mode was changed to 1 to return a report in Kraken format for further downstream analysis.

### Visualization

The kraken report for each sample obtained was uploaded onto the online version of the *shiny R* package [54]. The generated CSV tables and Sankey diagrams of all classified contigs generated for each sample were aggregated into a final HTML report.

## Results

Illumina sequencing of the libraries for each fauna species yielded over 5 million reads per species following adapter removal and quality control. The taxonomic analysis demonstrated a robust taxonomic breadth and resolution, with an average of 92% of organisms successfully classified at the species or genus level.

### Partitioning of food resources by Anopheles gambiae larvae and other fauna species

The *Anopheles* mosquito larvae shared nutritional resources not only with *Culex* species, which were the most abundant mosquito species identified, but also with other Diptera including *Toxorynchithes* and *Clunio* species (supplementary data 1). Specific instances of shared microbial and algal resources included *Stenotrophomonas maltophilia, Acinetobacter guillouiae, Pseudomonas putida*, and *Heterosigma akashiwo* with *Culex* species; *Goniomonas pacifica, Eutreptiella gymnastica, Chlamydomonas leiostraca*, and *Thalassiosira minuscula* with Toxorynchithes; as well as *Dunaliella tertiolecta, Paraphysomonas imperforata, Pseudomonas mendocina*, and *Brevundimonas diminuta* with *Clunio* species, (Fig 2).

**Fig 2.**
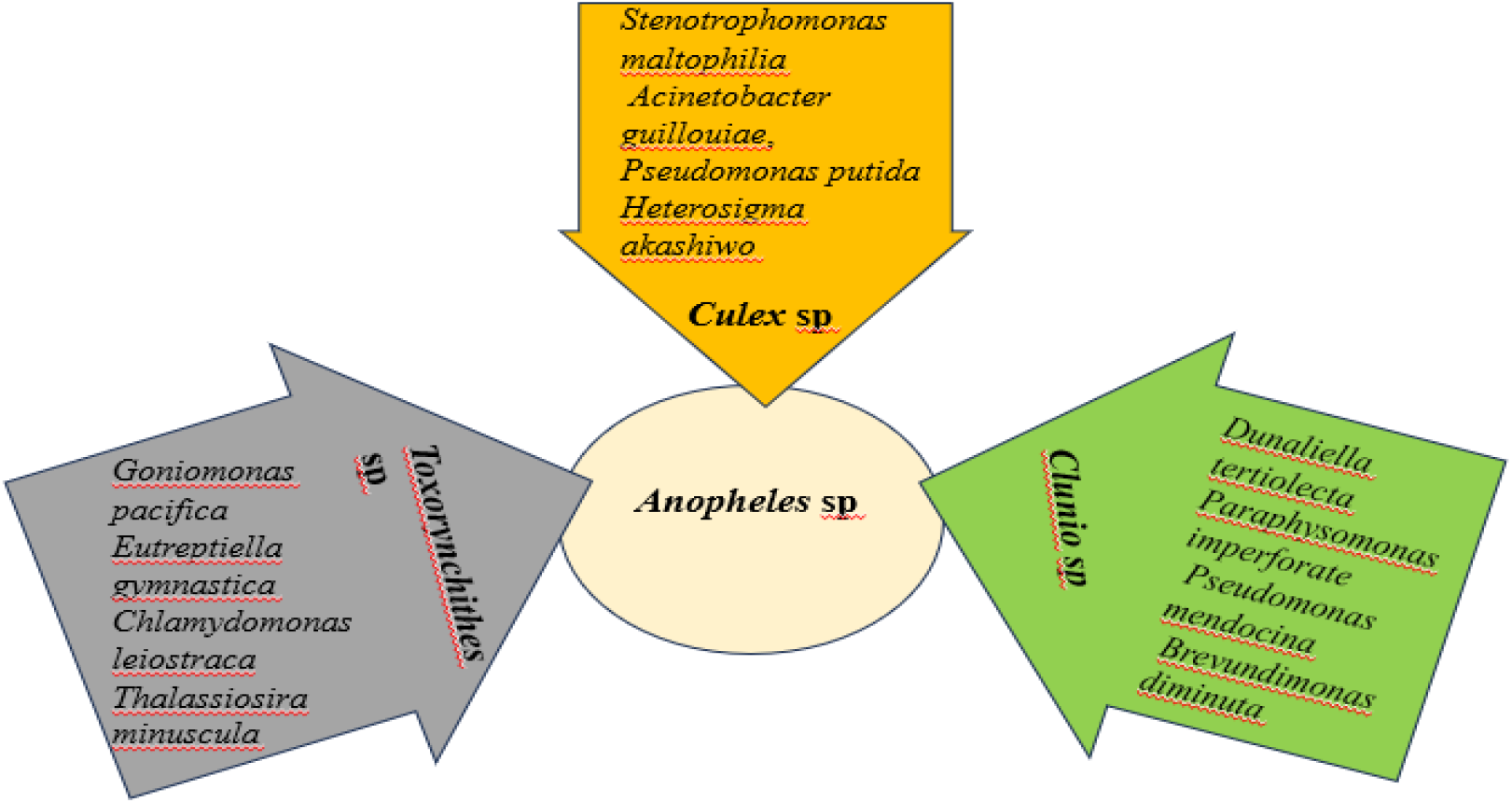
Some of the stomach compositions that *Anopheles* larvae share with the three Diptera species: (*Toxorynchithes* sp; *Culex* sp) and *Clunio* sp)

*Anopheles gambiae* larval diet was found to also exhibit dietary overlaps with non-Dipteran, detritus and filter-feeding organisms such as copepods and backswimmers. Specifically, they shared, *Rastonia pickettii Acinetobacter guillouiae, Heterosigma akashiwo, Goniomonas pacifica, Paraphysomonas imperforata, Chromonas cf. mesostigma, Diploscapter pachys*, and *Parastrongyloides trichosuri* with copepods (Fig 3) and *Stenotrophomonas maltophilia, Acinetobacter baumannii, Pseudomonas putida Diploscapter pachys, Goniomonas pacifica, Paraphysomonas imperforata, Chromonas cf. mesostigmata*, and *Trichuris trichiura* with backswimmer (Fig 3).

**Fig 3.**
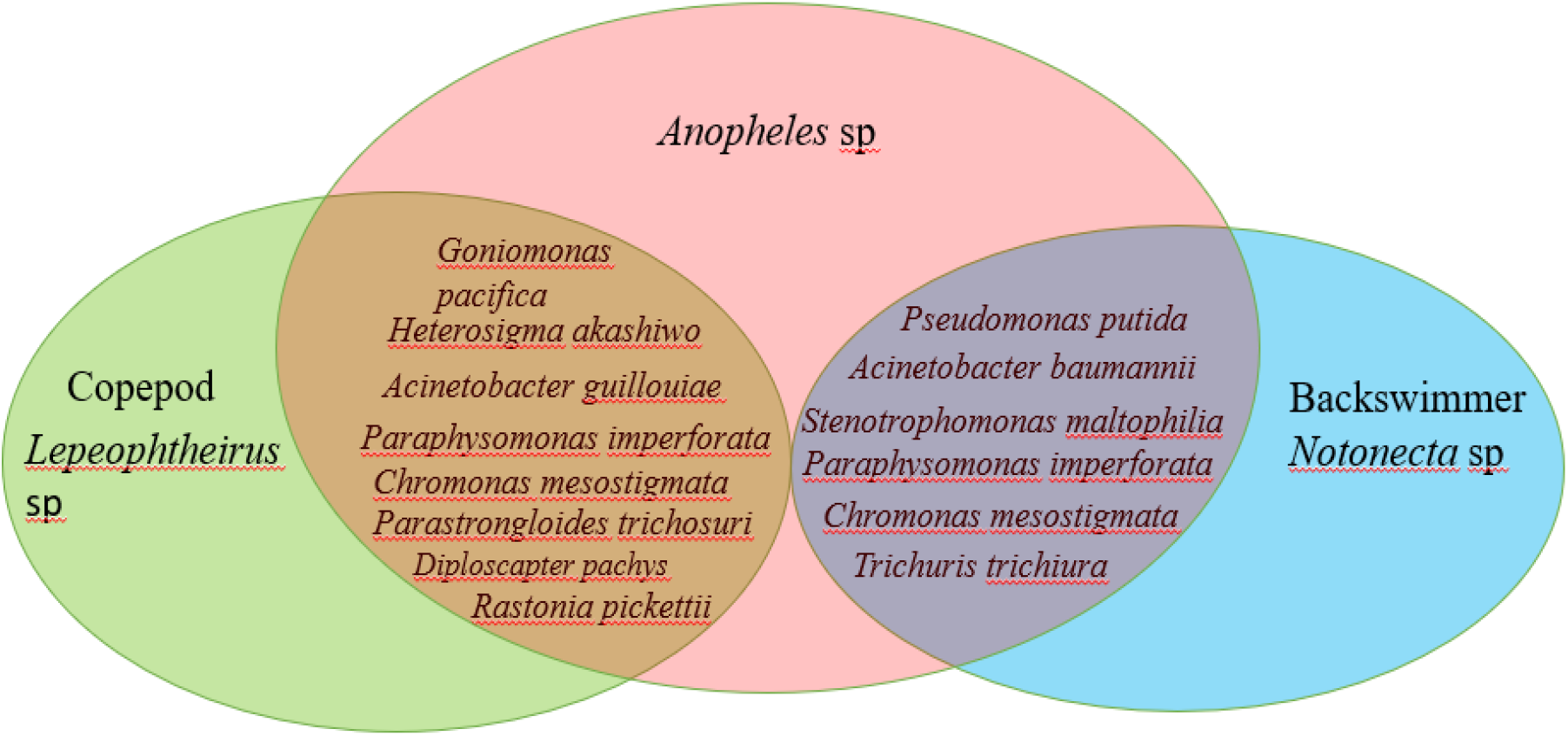
Some gut composition that *Anopheles* larvae share with non- Dipteran species: (*Lepeophtheirus sp*); (*Notonecta* sp)

### Relative abundances of Some Gut Composition of Anopheles larvae

The study found bacteria to be the major (70%) gut constituents of *Anopheles* larvae, in addition to protists, fungi, nematodes and plants (supplementary data 2). These included bacteria such as *Mycobacteroides abscessus*; fungi like *Trachydiscus minutus* and *Zancudomyces culisetae*; the filamentous bacterium *Hydrotalea sandarakina*, nematodes exemplified by *Brugia timori*; a variety of protozoans including *Phacus orbicularis, Euglena gracilis*, and *Euglenaria anabene*; and the alga *Chromera velia*; underscore the complexity of the *Anopheles* larvae diet, reflecting a wide range of ingested microorganisms and highlighting potential ecological interactions within their habitats. The relative abundance of some of the bacteria and algae found in *Anopheles* larvae gut are shown in Fig 4.

**Fig 4.**
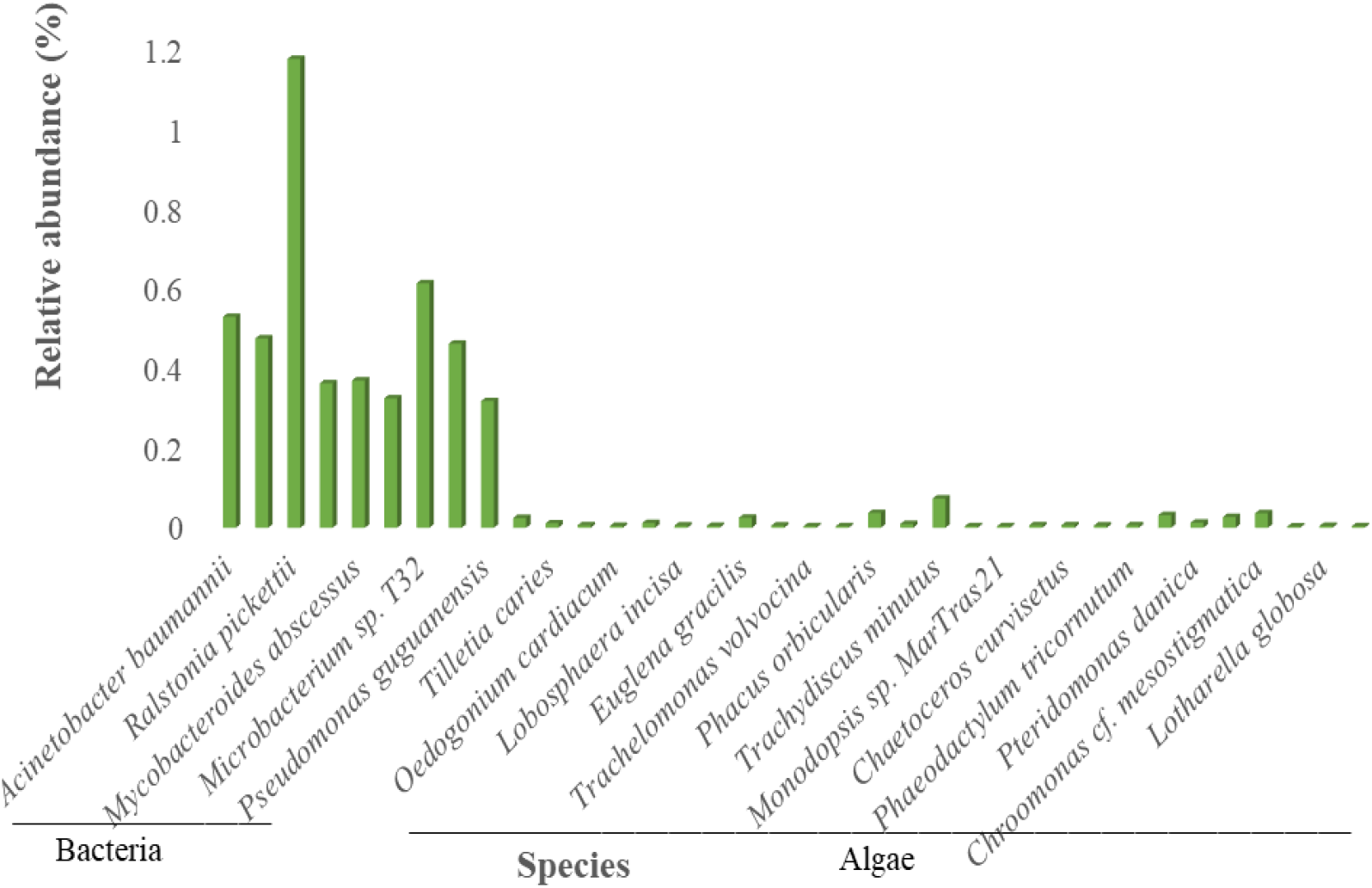
The relative abundance of some of the species found in *Anopheles* larvae’s gut

**Fig 4.**
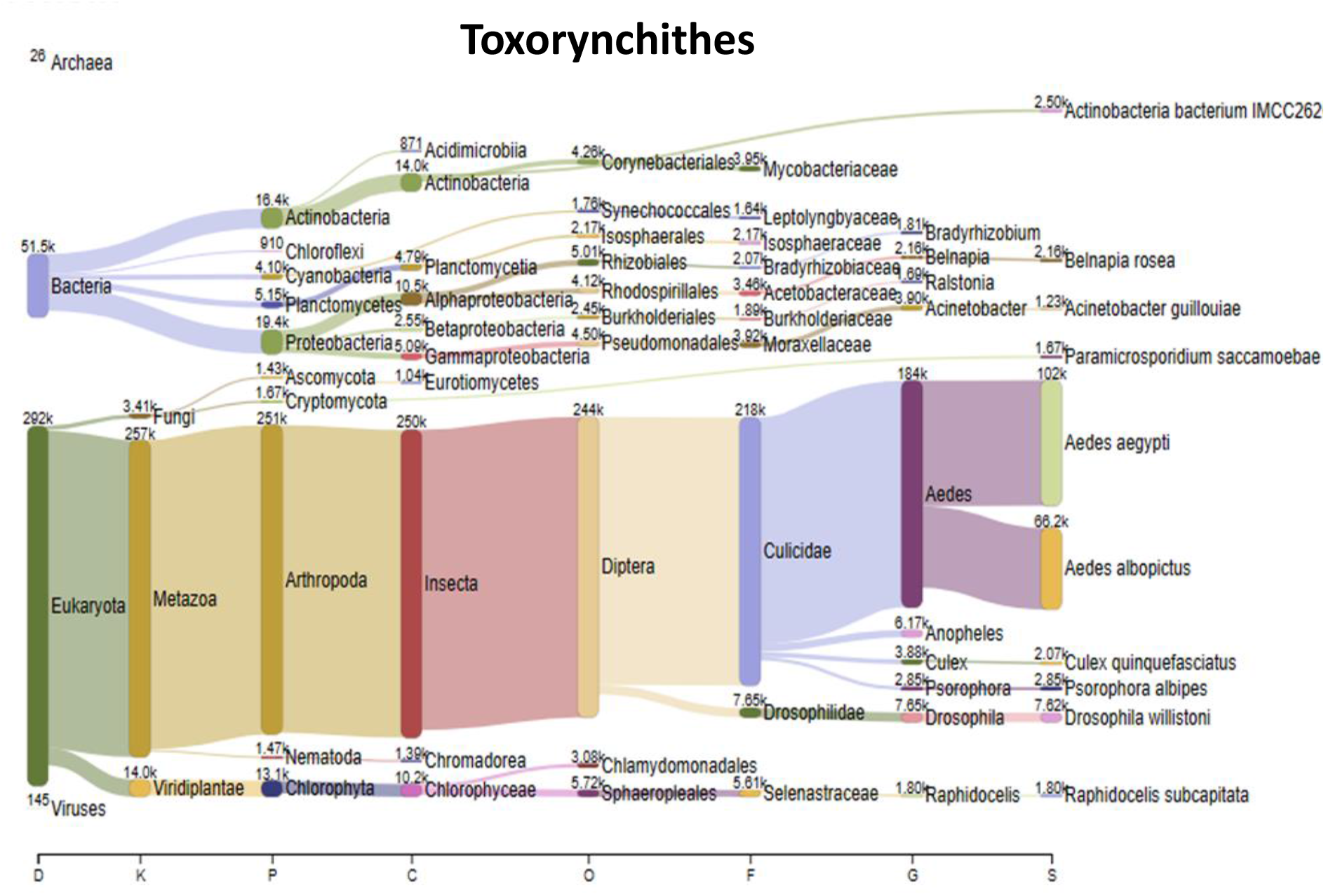
Taxonomic diet content distribution of Toxorynchithes larvae. The taxa are classified by their relative abundance in the Kingdom, Phylum, Genus and Species

### Diet composition of the selected predators of Anopheles gambiae larvae

This study analyzed the gut contents of aquatic fauna previously documented as predators of *Anopheles gambiae* larvae and identified within the study sites. The predatory species included *Toxorhynchites* spp., copepods, backswimmers (*Notonecta* spp.), damselflies, dragonflies, and mosquito fish (*Gambusia affinis*) [38-42].

Gut content analysis revealed a diverse range of macroinvertebrate families, including members of Culicidae, Reduviidae, Drosophilidae, Miridae, Cicadellidae, Aphididae, Formicidae, Lygaeidae, Liviidae, Clastopteridae, Taeniidae, Apidae, Glossinidae, Pteromalidae, Psychodidae, Calliphoridae, Chironomidae, and Halictidae. Detailed breakdowns of family-level composition and their relative proportions for each predator species are provided in Supplementary Data 3. In addition, the classification trees presented below (Figs 4-9) outline the complete taxonomic profiles of the prey identified in the guts of these predators.

The gut of *Toxorynchites* species showed a diet dominated by insects, with members of the Culicidae family, particularly *Aedes aegypti*, being the most significant dietary component (Fig. 4). Other Culicidae members identified within their diet were species of *Anopheles* and *Culex*.

The gut analysis of copepod backswimmer (*Notonecta* species) stomach contents highlighted a predominance of bacteria over eukaryotes (Figs 5 & 6) as well as some fungi and plants.

**Fig 5.**
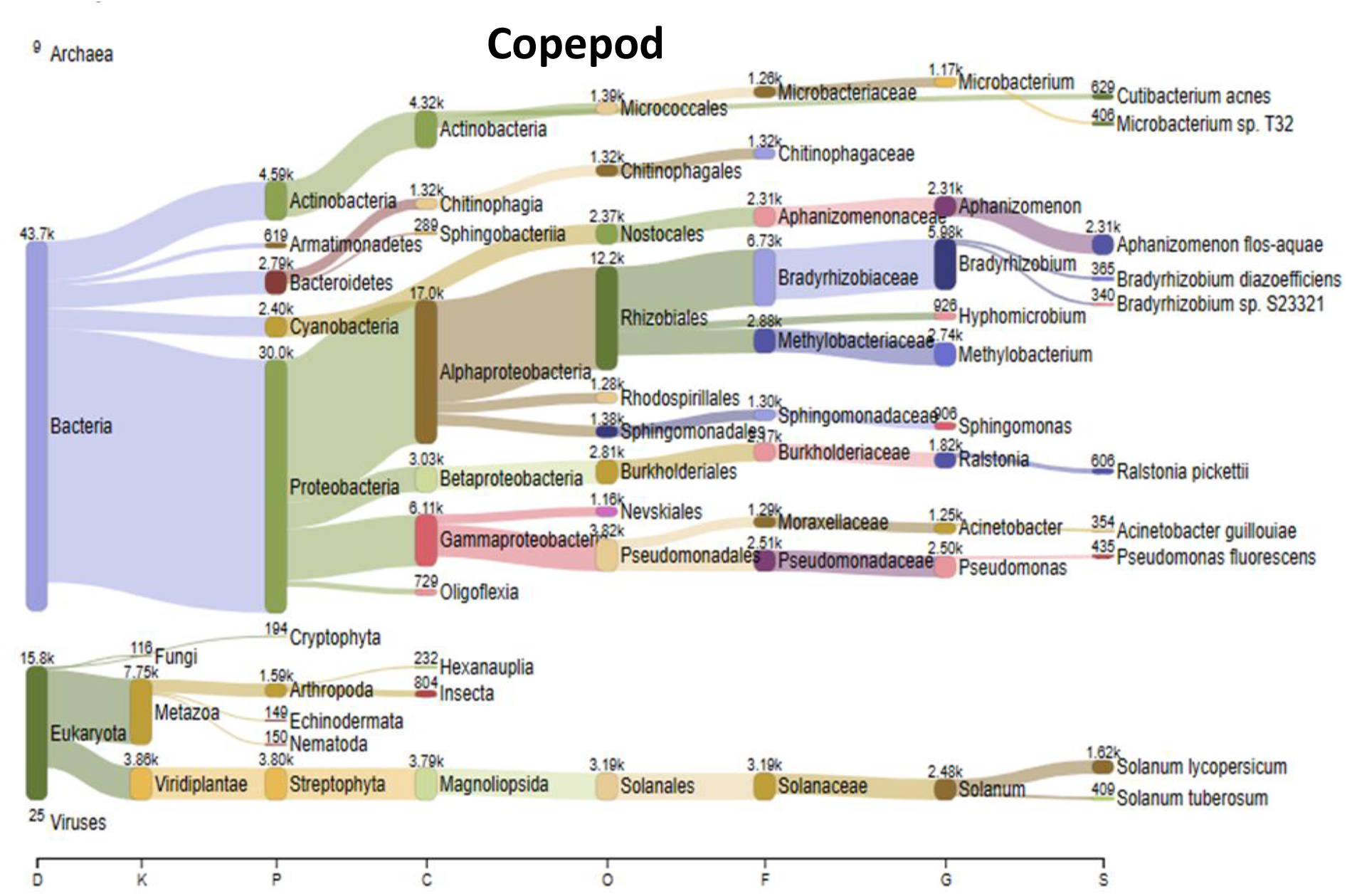
Taxonomic diet content distribution of Copepod. The taxa are classified by their relative abundance in the Kingdom, Phylum, Genus and Species

**Fig 6.**
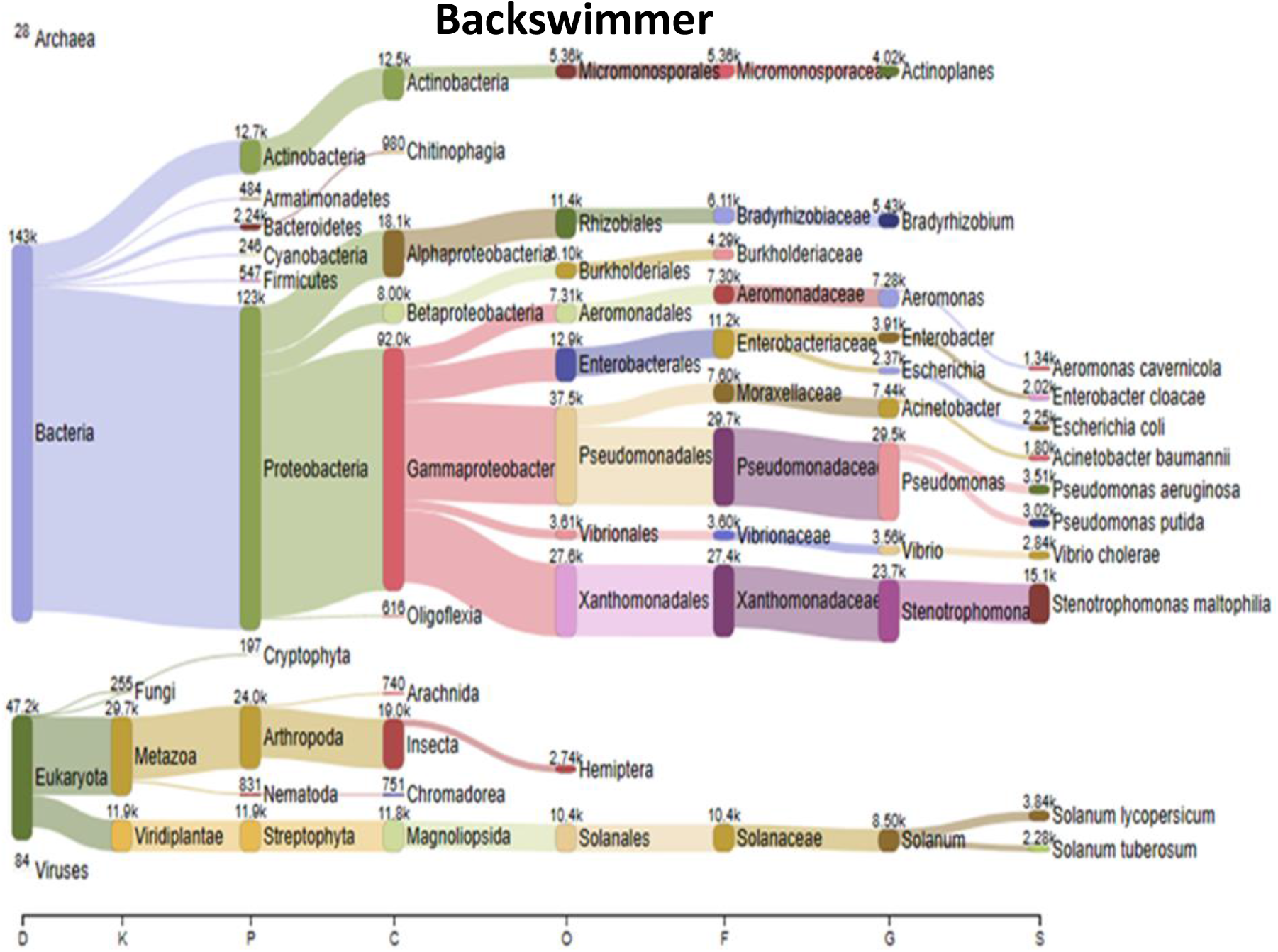
Taxonomic diet content distribution of Backswimmer (*Notonecta* sp). The taxa are classified by their relative abundance in the Kingdom, Phylum, Genus and Species

The gut contents of damselflies had a narrowed arthropod diet, which largely comprised culicine insects. Among these, *Culex quinquefasciatus* emerged as the most abundant species recorded. Additionally, traces of the *Anopheles* species were identified, accounting for a minor fraction (3.5%) of the total insect components consumed by damselflies (Fig. 7).

**Fig 7.**
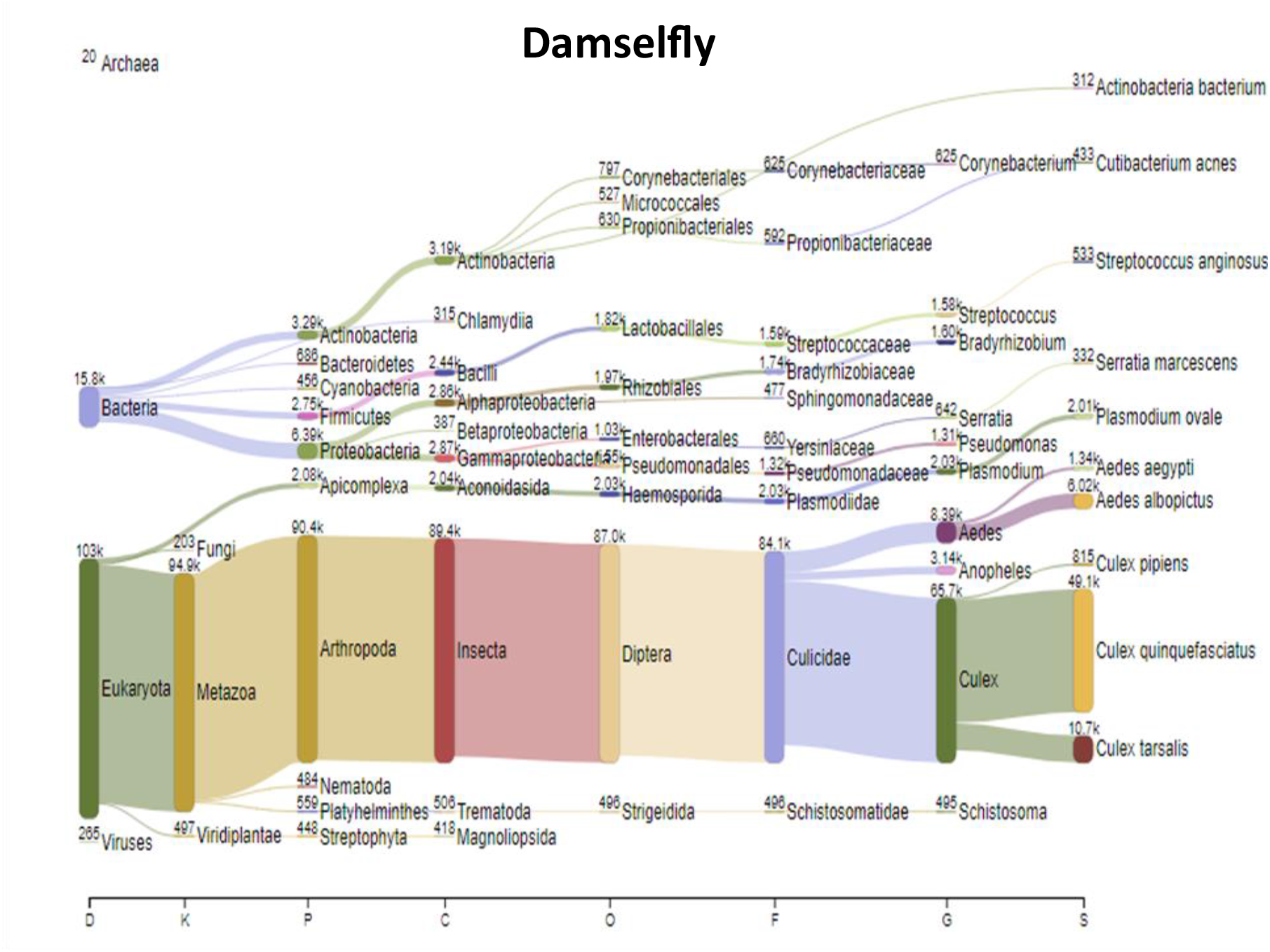
Taxonomic diet content distribution of Damselflies (*Ischnura* sp). The taxa are classified by their relative abundance in the Kingdom, Phylum, Genus and Species

Dragonflies on the other hand, had a more diverse arthropod diet than the damselfly. They also exhibited a dietary preference for other insect species other than *Anopheles* which represented only 1.9% of the total insect composition in the dragonfly diet (Fig. 8).

**Fig 8.**
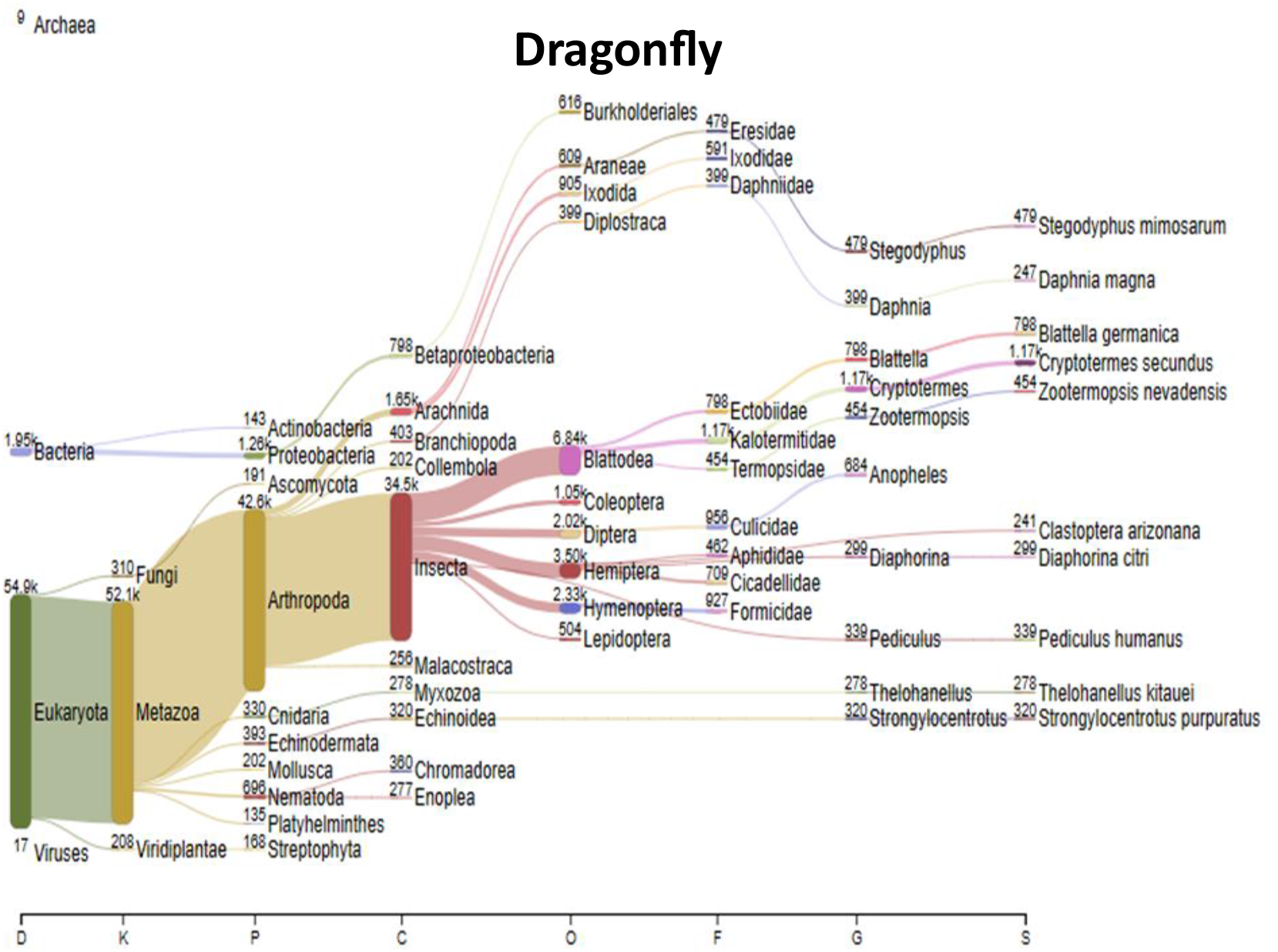
Taxonomic diet content distribution of Dragonfly (*Pantala* sp). The taxa are classified by their relative abundance in the Kingdom, Phylum, Genus and Species

The diet of *Gambusia affinis* primarily consisted of Platyhelminthes with a relatively minimal proportion of insects (Fig 9) including Diptera

**Fig 9.**
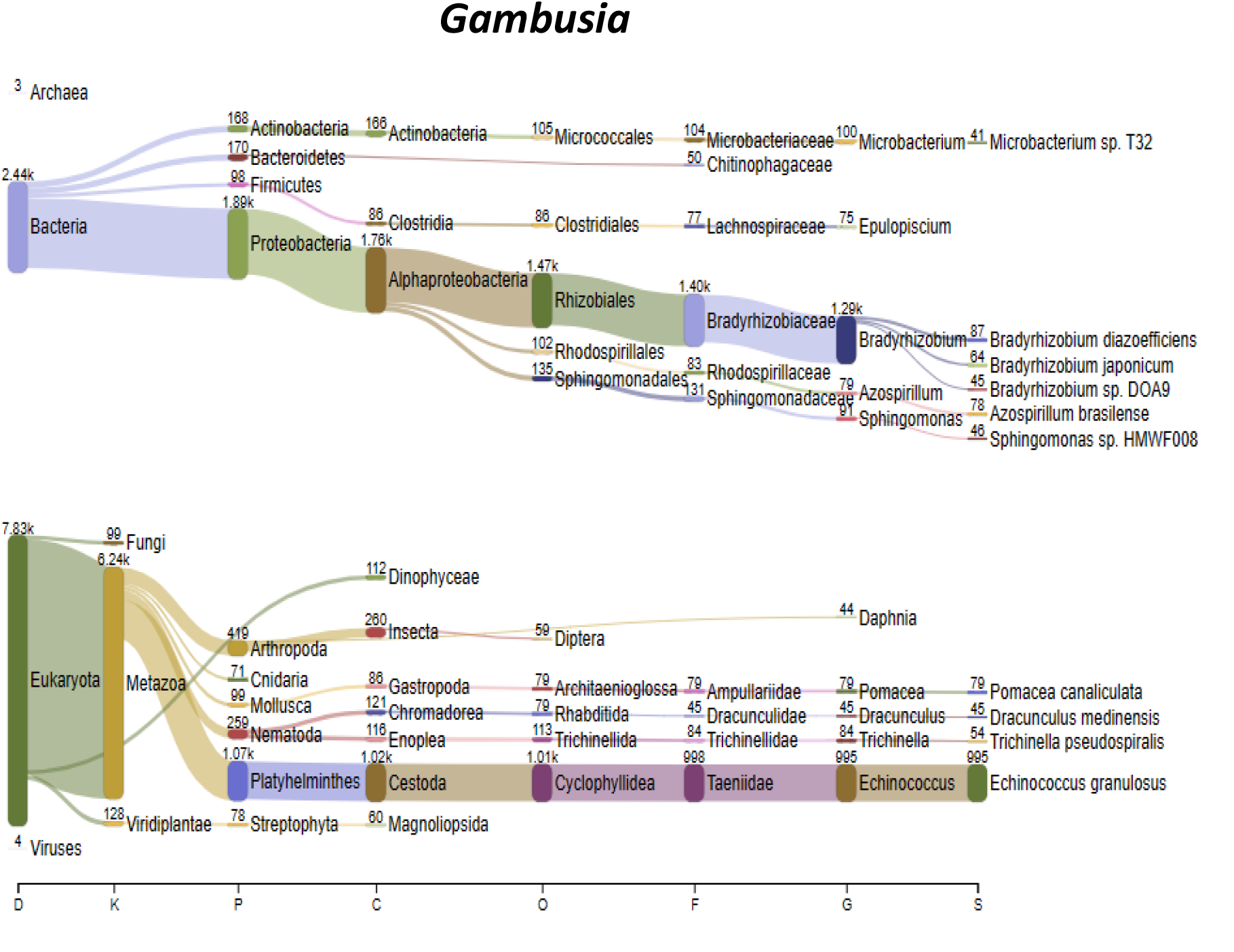
Taxonomic diet content distribution of *Gambusia affinis*. The taxa are classified by their relative abundance in the Kingdom, Phylum, Genus and Species

## Discussion

The use of shotgun metagenomic sequencing in this study provided a comprehensive analysis of the feeding ecology and trophic interactions involving *Anopheles gambiae* larvae and their predators. Previous research has established that the diet of *Anopheles* mosquitoes, primarily consists of bacteria, algae, diatoms, and detritus [54,55]. However, the present study uncovered additional dietary constituents, including nematodes and fungi, suggesting more intricate food constituents within the larval gut environment.

The dietary analysis also highlighted a complex web of trophic interactions, revealing both intra-and interspecific resource sharing within the aquatic ecosystem. Notably, *An. gambiae* larvae were found to share dietary resources not only with other Diptera species but also with copepods and backswimmers. It is believed that these interactions occur despite the predatory tendencies of copepods and backswimmers, as these organisms exhibit detritus and filter-feeding behaviors [56] and can coexist with *Anopheles* larvae, probably due to resource partitioning strategies.

This discovery has important implications for malaria control strategies, particularly in the context of gene drive applications. One critical concern with gene drive interventions (particularly those aimed at *Anopheles* reduction/loss) is the potential ecological risk of conferring a competitive edge to other mosquito vectors. However, the observed food partitioning behavior among *An. gambiae* larvae and other detritus and filter feeders reduce the likelihood of such competitive advantage. In the event of population reduction through gene drive, resource-sharing mechanisms among the mosquitoes and the other detritus feeders could help prevent other vector species from gaining a significant advantage, thereby mitigating unintended ecological consequences.

In ecological studies, predator species are typically classified as either specialists or generalists based on their dietary habits. Specialists (monophagic) focus on a single prey type, while generalists (polyphagic) consume a variety of prey species [57, 58]. Contrary to expectations, none of the predator species employed in this study exhibited exclusive predation on *Anopheles* mosquito larvae. Instead, they predominantly consumed other prey, demonstrating a generalist feeding behavior. This result again gives important implications for gene drive application, as the absence of specialist predators targeting *Anopheles* larvae suggests minimal risk of ecosystem disruption through predator loss.

Previous studies [38], have documented mosquito predators in larval habitats, including backswimmers (*Notonecta* ), dragonflies and damselflies (Odonata), Toxorhynchites mosquitoes [39], chironomids [40], copepods (various species) [41], and mosquito fish (*Gambusia affinis*) [42]. In this study, our analysis did not detect *Anopheles* DNA in the gut content of *G. affinis*. although it has been proposed as a key predator for *Anopheles* larvae in larval source management strategies [42], Our observation could be in agreement with studies conducted in natural water bodies, which reported that mosquito larvae do not form a substantial part of the *G. affinis* diet [59]. However, there is also the likelihood that the absence of *Anopheles* DNA could also be due to small sample size, since only a single *G. affinis* specimen was analyzed (Due to inadequate collected *G. affinis* samples). A similar limitation applies to the copepod sample, which was also assessed as a single sample. We therefore propose further research with larger sample sizes to draw more definitive conclusions about the dietary habits of *G. affinis* and copepods in natural settings.

The findings of this study diverge from existing literature suggesting that Odonata, particularly dragonflies, prey significantly on mosquito larvae [60]. *Anopheles* larvae were found to constitute only a small fraction (1.22%) of the arthropod composition in the natural diets of these predators. This suggests that Odonata species may primarily consume *Anopheles* larvae only under certain conditions, such as in laboratory settings or environments lacking alternative prey [61]. In a similar pattern, we found *Anopheles* as a minute (2.5%) part of *Toxorynchithes* species’ diet but the results corroborate previous reports identifying *Toxorhynchites* mosquitoes as significant predators of *Aedes aegypti* larvae [62, 39]. The limited predation on *Anopheles* larvae may align with predictions from optimal foraging theory, which indicates that prey size and profitability influence predator preferences [3, 59]. As *Anopheles* larvae are relatively small, they may not offer sufficient energetic returns to be favored in the natural diets of these predators.

This study is meaningful for informing the design of effective mosquito control strategies, particularly those involving biological control and gene drive applications.

## Conclusion

This study has uncovered complex trophic interactions involving *Anopheles gambiae* larvae and other organisms within freshwater ecosystems. The absence of exclusive predation on *An. gambiae* larvae and competitive edge by another mosquito vector suggest that vector control strategies focused on reducing mosquito larvae are unlikely to disrupt ecological balance. These insights have important implications for the implementation of *Anopheles* mosquito control strategies, such as gene drive technology.

Additionally, the it has provided valuable information on the aquatic fauna of the study area, which can serve as a baseline for developing a macroinvertebrate identification database for freshwater systems in Ghana.

A complementary study that further explores the ecological role of *An. gambiae* larvae in these freshwater habitats is presented in a subsequent publication.

## Limitation

Some fauna bore exotic names in the taxonomic classification dendrograms of the results. For instance, we found termite species in the predatory guts being classified as *Cryptotermes and zootermopsis* species instead of *Macrotermes* or other African species. This discrepancy is perhaps a consequence of Ghana’s lack of a comprehensive aquatic database, leading to species found in Ghana being assigned foreign names. Further research is being undertaken to elucidate these observations.

## Acknowledgments

We are extremely grateful to Benard Adams, Collins Awuah, Freeman Awitor, Richmond Gave, and Seraphim N.A. Tetteh for their technical assistance on the field and in the laboratory.

