## supplementary data 1 for "Trophic Interactions of *Anopheles Gambiae* Mosquito Larvae in Aquatic Ecosystem: A Metagenomics Approach"

*Classification tree showing the gut constituents of Anopheles and other dipteran (excluding Toxorynchithes sp) larvae*

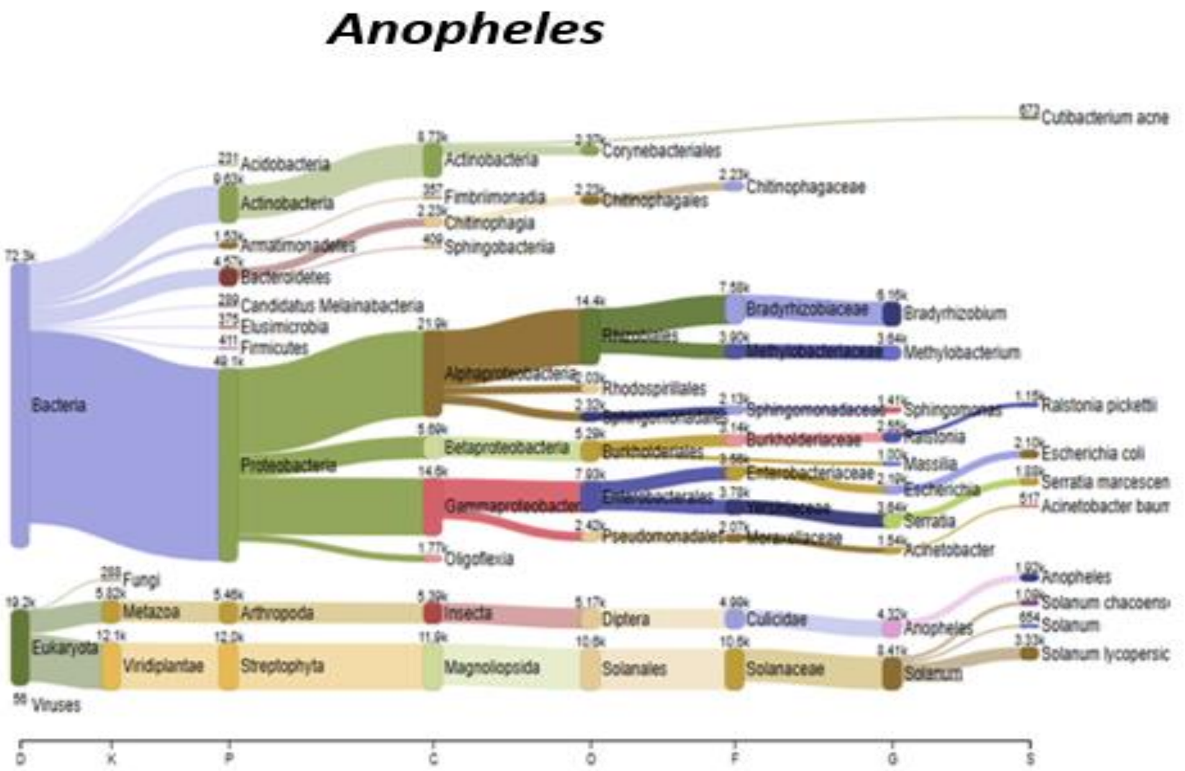


A) Taxonomic diet content distribution of *Anopheles* larvae. The taxa are classified by their relative abundance according to Kingdom, Phylum, Genus and Species.

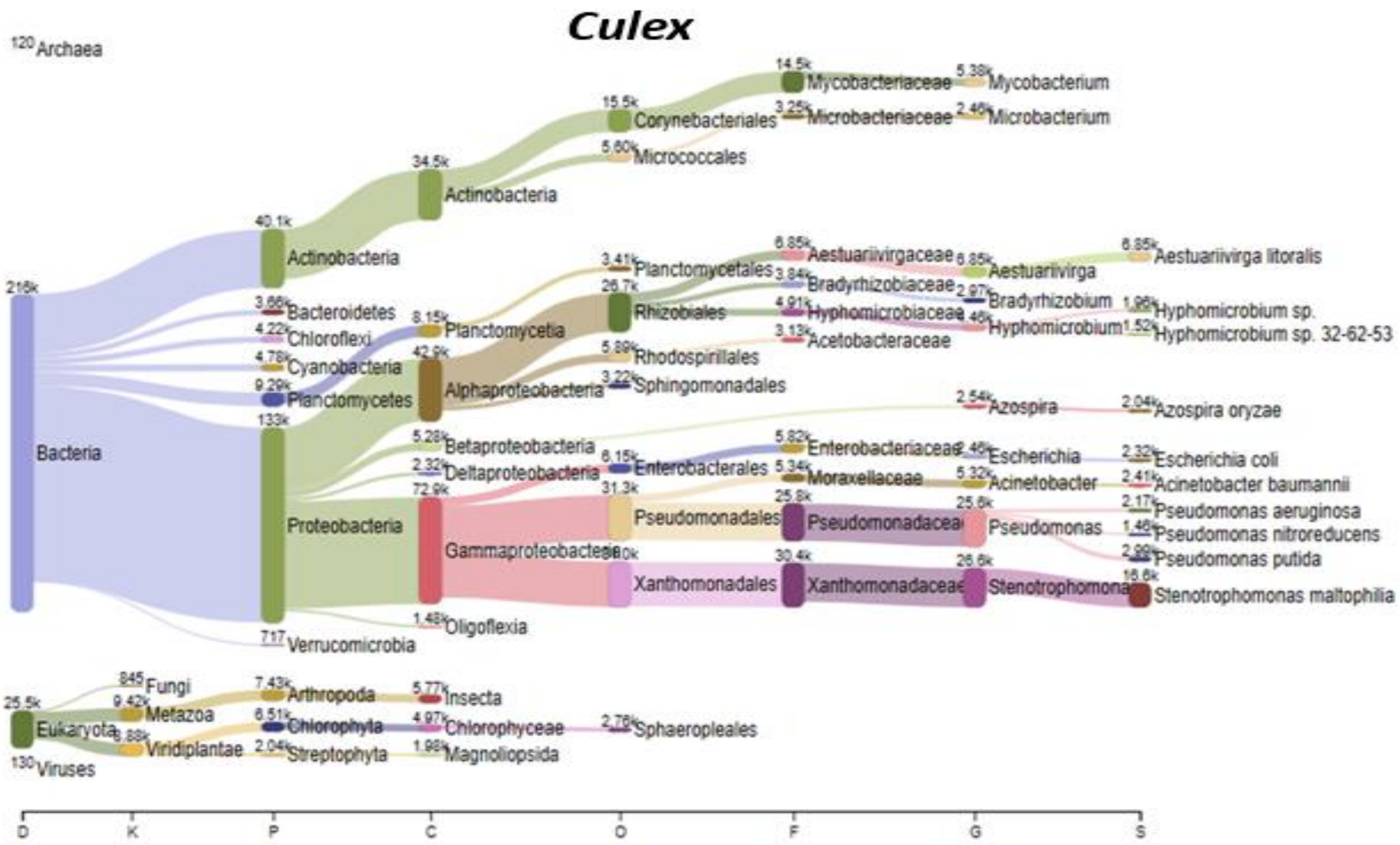


b) Taxonomic diet content distribution of *Culex* larvae. The taxa are classified by their relative abundance according to Kingdom, Phylum, Genus and Species

**Clunio**

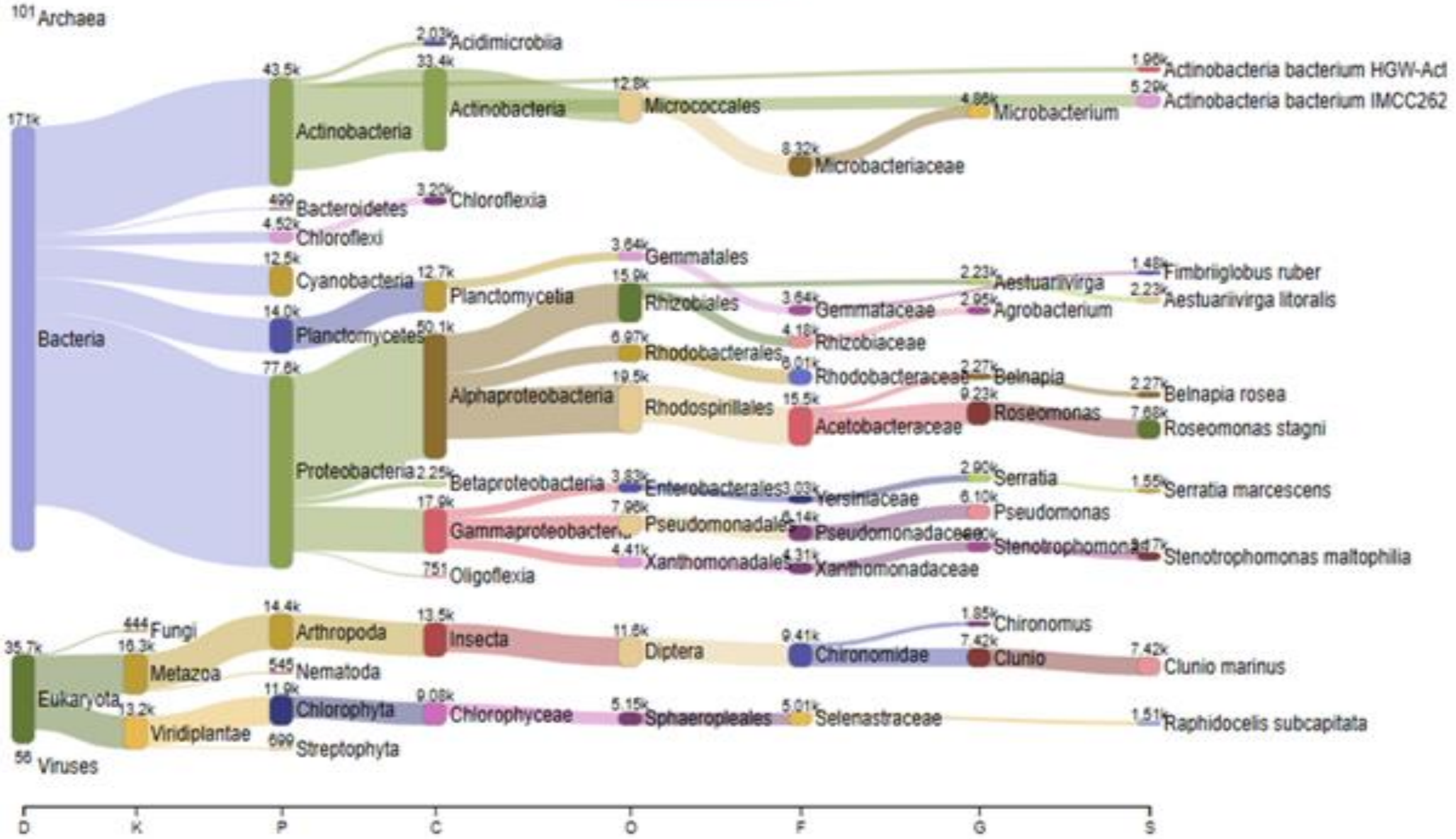


**c)**Taxonomic diet content distribution of Chironomid larvae. The taxa are classified by their relative abundance according to Kingdom, Phylum, Genus and Species
