## supplementary data 2 for "Trophic Interactions of *Anopheles Gambiae* Mosquito Larvae in Aquatic Ecosystem: A Metagenomics Approach"

*Relative abundance of gut constituents of Anopheles larvae*

| Percentage | CladeRea | TaxRank | TaxID | *Name* | Depth | TaxLineage |
|---|---|---|---|---|---|---|
| **96.5672** | **93479** | **-** | **1** | **-_root** | **0** | **-_root** |
| **96.4176** | **93333** | **-** | **131567** | **-_cellular organisms** | **1** | **-_root\|-_cellular organisms** |
| **0.295115** | **288** | **K** | **4751** | **k_Fungi** | **4** | **-_root\|-_cellular organisms\|d_Eukaryota\|-_Opisthokonta\|** |
| 0.0102558 | 13 | S | 93612 | *s_Exserohilum turcicum* | 17 | -_root\|-_cellular organisms\|d_Eukaryota\|-_Opisthokonta\| |
| 0.0110447 | 14 | S | 13290 | *s_Tilletia caries* | 12 | -_root\|-_cellular organisms\|d_Eukaryota\|-_Opisthokonta\| |
| 0.00631124 | 8 | S | 43049 | *s_Tilletia indica* | 12 | -_root\|-_cellular organisms\|d_Eukaryota\|-_Opisthokonta\| |
| 0.00512353 | 5 | S | 1708541 | *s_Wallemia mellicola* | 12 | -_root\|-_cellular organisms\|d_Eukaryota\|-_Opisthokonta\| |
| 0.00409882 | 4 | S | 396024 | *s_Aspergillus ruber* | 15 | -_root\|-_cellular organisms\|d_Eukaryota\|-_Opisthokonta\| |
| 0.0143459 | 14 | S | 45151 | *s_Pyrenophora tritici-repentis* | 17 | -_root\|-_cellular organisms\|d_Eukaryota\|-_Opisthokonta\| |
| 0.0245929 | 24 | S | 1213189 | *s_Zancudomyces culisetae* | 12 | -_root\|-_cellular organisms\|d_Eukaryota\|-_Opisthokonta\| |
| **12.3549** | **12057** | **K** | **33090** | **k_Viridiplantae (including algae )** | **3** | **-_root\|-_cellular organisms\|d_Eukaryota\|k_Viridiplantae** |
| 3.41125 | 3329 | S | 4081 | *s_Solanum lycopersicum* | 23 | -_root\|-_cellular organisms\|d_Eukaryota\|k_Viridiplantae\| |
| 1.11386 | 1087 | S | 4108 | *s_Solanum chacoense* | 22 | -_root\|-_cellular organisms\|d_Eukaryota\|k_Viridiplantae\| |
| 0.212114 | 207 | S | 4072 | *s_Capsicum annuum* | 22 | -_root\|-_cellular organisms\|d_Eukaryota\|k_Viridiplantae\| |
| 0.264374 | 258 | S | 4097 | *s_Nicotiana tabacum* | 22 | -_root\|-_cellular organisms\|d_Eukaryota\|k_Viridiplantae\| |
| 0.0256176 | 25 | S | 4530 | *s_Oryza sativa* | 22 | -_root\|-_cellular organisms\|d_Eukaryota\|k_Viridiplantae\| |
| 0.0646902 | 82 | S | 1034604 | *s_Chlamydomonas leiostraca* | 10 | -_root\|-_cellular organisms\|d_Eukaryota\|k_Viridiplantae\| |
| 0.0213004 | 27 | S | 225041 | *s_Chlamydomonas chlamydogam* | 10 | -_root\|-_cellular organisms\|d_Eukaryota\|k_Viridiplantae\| |
| 0.0102558 | 13 | S | 1486919 | *s_Chlamydomonas euryale* | 10 | -_root\|-_cellular organisms\|d_Eukaryota\|k_Viridiplantae\| |
| 0.00788905 | 10 | S | 1157962 | *s_Chlamydomonas eustigma* | 10 | -_root\|-_cellular organisms\|d_Eukaryota\|k_Viridiplantae\| |
| 0.00473343 | 6 | S | 3055 | *s_Chlamydomonas reinhardtii* | 10 | -_root\|-_cellular organisms\|d_Eukaryota\|k_Viridiplantae\| |
| 0.0339229 | 43 | S | 3047 | *s_Dunaliella tertiolecta* | 10 | -_root\|-_cellular organisms\|d_Eukaryota\|k_Viridiplantae\| |
| 0.00552233 | 7 | S | 312850 | *s_Lobosphaera incisa* | 10 | -_root\|-_cellular organisms\|d_Eukaryota\|k_Viridiplantae\| |
| 0.00473343 | 6 | S | 3165 | *s_Tetraselmis striata* | 10 | -_root\|-_cellular organisms\|d_Eukaryota\|k_Viridiplantae\| |
| 0.00473343 | 6 | S | 156128 | *s_Nephroselmis pyriformis* | 9 | -_root\|-_cellular organisms\|d_Eukaryota\|k_Viridiplantae\| |
| 0.025245 | 32 | S | 3039 | *s_Euglena gracilis* | 10 | -_root\|-_cellular organisms\|d_Eukaryota\|p_Euglenozoa\|-_ |
| 0.262705 | 333 | S | 2829 | *s_Heterosigma akashiwo* | 9 | -_root\|-_cellular organisms\|d_Eukaryota\|-_Stramenopiles |
| 0.0733682 | 93 | S | 1032745 | *s_Trachydiscus minutus* | 8 | -_root\|-_cellular organisms\|d_Eukaryota\|-_Stramenopiles |
| 0.0157781 | 20 | S | 385413 | *s_Thalassiosira minuscula* | 11 | -_root\|-_cellular organisms\|d_Eukaryota\|-_Stramenopiles |
| 0.00631124 | 8 | S | 230516 | *s_Chaetoceros curvisetus* | 11 | -_root\|-_cellular organisms\|d_Eukaryota\|-_Stramenopiles |
| 0.0315562 | 40 | S | 37642 | *s_Paraphysomonas imperforata* | 9 | -_root\|-_cellular organisms\|d_Eukaryota\|-_Stramenopiles |
| 0.0126225 | 16 | S | 38822 | *s_Pteridomonas danica* | 8 | -_root\|-_cellular organisms\|d_Eukaryota\|-_Stramenopiles |
| 0.148314 | 188 | S | 195067 | *s_Goniomonas pacifica* | 7 | -_root\|-_cellular organisms\|d_Eukaryota\|p_Cryptophyta\|c |
| 0.0268228 | 34 | S | 464988 | *s_Hemiselmis andersenii* | 7 | -_root\|-_cellular organisms\|d_Eukaryota\|p_Cryptophyta\|c |
| 0.0362896 | 46 | S | 478117 | *s_Chroomonas cf. mesostigmatica* | 7 | -_root\|-_cellular organisms\|d_Eukaryota\|p_Cryptophyta\|o_Pyrer |
| 0.00473343 | 6 | S | 91324 | *s_Lotharella globosa* | 7 | -_root\|-_cellular organisms\|d_Eukaryota\|-_Rhizaria\|p_Cercozoa |
| **0.184447** | **180** | **P** | **6231** | **p_Nematoda** | **9** | **-_root\|-_cellular organisms\|d_Eukaryota\|-_Opisthokonta\|** |
| 0.0768529 | 75 | S | 2018661 | *s_Diploscapter pachys* | 18 | -_root\|-_cellular organisms\|d_Eukaryota\|-_Opisthokonta\| |
| 0.00614823 | 6 | S | 31234 | *s_Caenorhabditis remanei* | 18 | -_root\|-_cellular organisms\|d_Eukaryota\|-_Opisthokonta\| |
| 0.045087 | 44 | S | 131310 | *s_Parastrongyloides trichosuri* | 17 | -_root\|-_cellular organisms\|d_Eukaryota\|-_Opisthokonta\| |
| 0.00922235 | 9 | S | 42155 | *s_Brugia timori* | 17 | -_root\|-_cellular organisms\|d_Eukaryota\|-_Opisthokonta\| |
| 0.0153706 | 15 | S | 36087 | *s_Trichuris trichiura* | 15 | -_root\|-_cellular organisms\|d_Eukaryota\|-_Opisthokonta\| |
| 0.00409882 | 4 | S | 990121 | *s_Trichinella patagoniensis* | 15 | -_root\|-_cellular organisms\|d_Eukaryota\|-_Opisthokonta\| |
| **0.1742** | **170** | **-** | **33630** | **-_Alveolata** | **3** | **-_root\|-_cellular organisms\|d_Eukaryota\|-_Alveolata** |
| 0.0796794 | 101 | S | 5911 | *s_Tetrahymena thermophila* | 11 | -_root\|-_cellular organisms\|d_Eukaryota\|-_Alveolata\|p_Ciliophora |
| 0.0236671 | 30 | S | 5932 | *s_Ichthyophthirius multifiliis* | 10 | -_root\|-_cellular organisms\|d_Eukaryota\|-_Alveolata\|p_Ci |
| 0.0804683 | 102 | S | 5888 | *s_Paramecium tetraurelia* | 10 | -_root\|-_cellular organisms\|d_Eukaryota\|-_Alveolata\|p_Ci |
| 0.0291895 | 37 | S | 266149 | *s_Pseudocohnilembus persalinus* | 11 | -_root\|-_cellular organisms\|d_Eukaryota\|-_Alveolata\|p_Ci |
| 0.00788905 | 10 | S | 1172189 | *s_Oxytricha trifallax* | 12 | -_root\|-_cellular organisms\|d_Eukaryota\|-_Alveolata\|p_Ciliophora |
| 0.00710014 | 9 | S | 5963 | *s_Stentor coeruleus* | 10 | -_root\|-_cellular organisms\|d_Eukaryota\|-_Alveolata\|p_Ciliophora |
| 0.119125 | 151 | S | 160619 | *s_Kryptoperidinium foliaceum* | 8 | -_root\|-_cellular organisms\|d_Eukaryota\|-_Alveolata\|c_Di |
| 0.0173559 | 22 | S | 2951 | *s_Symbiodinium microadriaticum* | 8 | -_root\|-_cellular organisms\|d_Eukaryota\|-_Alveolata\|c_Di |

| | | | | | | |
|---|---|---|---|---|---|---|
| **74.0524** | **72267** | **D** | **2** | **d_Bacteria** | **2** | **-_root\|-_cellular organisms\|d_Bacteria** |
| 0.543555 | 689 | S | 300 | s_*Pseudomonas mendocina* | 9 | -_root\|-_cellular organisms\|d_Bacteria\|p_Proteobacteria\| |
| 0.36053 | 457 | S | 301 | s_*Pseudomonas oleovorans* | 10 | -_root\|-_cellular organisms\|d_Bacteria\|p_Proteobacteria\| |
| 0.220893 | 280 | S | 287 | s_*Pseudomonas aeruginosa* | 9 | -_root\|-_cellular organisms\|d_Bacteria\|p_Proteobacteria\| |
| 0.300573 | 381 | S | 303 | s_*Pseudomonas putida* | 9 | -_root\|-_cellular organisms\|d_Bacteria\|p_Proteobacteria\| |
| 0.317929 | 403 | S | 1198456 | s_*Pseudomonas guguanensis* | 8 | -_root\|-_cellular organisms\|d_Bacteria\|p_Proteobacteria\| |
| 0.106502 | 135 | S | 101564 | s_*Pseudomonas alcaliphila* | 8 | -_root\|-_cellular organisms\|d_Bacteria\|p_Proteobacteria\| |
| 0.0804683 | 102 | S | 1274359 | s_*Pseudomonas sihuiensis* | 8 | -_root\|-_cellular organisms\|d_Bacteria\|p_Proteobacteria\| |
| 1.0098 | 1280 | S | 106649 | s_*Acinetobacter guillouiae* | 8 | -_root\|-_cellular organisms\|d_Bacteria\|p_Proteobacteria\| |
| 0.336073 | 426 | S | 106648 | s_*Acinetobacter bereziniae* | 8 | -_root\|-_cellular organisms\|d_Bacteria\|p_Proteobacteria\| |
| 0.529773 | 517 | S | 470 | s_*Acinetobacter baumannii* | 9 | -_root\|-_cellular organisms\|d_Bacteria\|p_Proteobacteria\| |
| 2.23181 | 2829 | S | 40324 | s_*Stenotrophomonas maltophilia* | 9 | -_root\|-_cellular organisms\|d_Bacteria\|p_Proteobacteria\| |
| 1.92235 | 1876 | S | 615 | s_*Serratia marcescens* | 8 | -_root\|-_cellular organisms\|d_Bacteria\|p_Proteobacteria\| |
| 0.110447 | 140 | S | 82996 | s_*Serratia plymuthica* | 8 | -_root\|-_cellular organisms\|d_Bacteria\|p_Proteobacteria\| |
| 1.29459 | 1641 | S | 358 | s_*Agrobacterium tumefaciens* | 10 | -_root\|-_cellular organisms\|d_Bacteria\|p_Proteobacteria\| |
| 0.123858 | 157 | S | 1183412 | s_*Agrobacterium deltaense* | 9 | -_root\|-_cellular organisms\|d_Bacteria\|p_Proteobacteria\| |
| 0.825194 | 1046 | S | 293 | s_*Brevundimonas diminuta* | 8 | -_root\|-_cellular organisms\|d_Bacteria\|p_Proteobacteria\| |
| 0.23036 | 292 | S | 2056845 | s_*Sphingomonas sp. HMWF008* | 9 | -_root\|-_cellular organisms\|d_Bacteria\|p_Proteobacteria\| |
| 0.155414 | 197 | S | 1296669 | s_*Aquabacterium olei* | 9 | -_root\|-_cellular organisms\|d_Bacteria\|p_Proteobacteria\| |
| 1.17739 | 1149 | S | 329 | s_*Ralstonia pickettii* | 8 | -_root\|-_cellular organisms\|d_Bacteria\|p_Proteobacteria\| |
| 0.116758 | 148 | S | 305 | s_*Ralstonia solanacearum* | 8 | -_root\|-_cellular organisms\|d_Bacteria\|p_Proteobacteria\| |
| 0.116758 | 148 | S | 190721 | s_*Ralstonia insidiosa* | 8 | -_root\|-_cellular organisms\|d_Bacteria\|p_Proteobacteria\| |
| 0.613768 | 778 | S | 1776083 | s_*Microbacterium sp. T32* | 10 | -_root\|-_cellular organisms\|d_Bacteria\|-_Terrabacteria gr |
| 0.104135 | 132 | S | 2033 | s_*Microbacterium testaceum* | 9 | -_root\|-_cellular organisms\|d_Bacteria\|-_Terrabacteria gr |
| 0.0930908 | 118 | S | 113567 | s_*Actinoplanes italicus* | 9 | -_root\|-_cellular organisms\|d_Bacteria\|-_Terrabacteria gr |
| 0.0836239 | 106 | S | 113562 | s_*Actinoplanes derwentensis* | 9 | -_root\|-_cellular organisms\|d_Bacteria\|-_Terrabacteria gr |
| 0.082835 | 105 | S | 1869 | s_*Actinoplanes utahensis* | 9 | -_root\|-_cellular organisms\|d_Bacteria\|-_Terrabacteria gr |
| 0.689627 | 673 | S | 1747 | s_*Cutibacterium acnes* | 9 | -_root\|-_cellular organisms\|d_Bacteria\|-_Terrabacteria gr |
| 0.250028 | 244 | S | 1355477 | s_*Bradyrhizobium diazoefficiens* | 8 | -_root\|-_cellular organisms\|d_Bacteria\|p_Proteobacteria\| |
| 0.317659 | 310 | S | 409 | s_*Methylobacterium sp.* | 9 | -_root\|-_cellular organisms\|d_Bacteria\|p_Proteobacteria\| |
| 0.462142 | 451 | S | 39956 | s_*Methylobacterium mesophilicur* | 8 | -_root\|-_cellular organisms\|d_Bacteria\|p_Proteobacteria\| |
| 0.475463 | 464 | S | 34062 | s_*Moraxella osloensis* | 8 | -_root\|-_cellular organisms\|d_Bacteria\|p_Proteobacteria\| |
| 0.270522 | 264 | S | 1805166 | s_*Gammaproteobacteria bacteriu* | 7 | -_root\|-_cellular organisms\|d_Bacteria\|p_Proteobacteria\| |
| 0.362746 | 354 | S | 1977087 | s_*Proteobacteria bacterium* | 6 | -_root\|-_cellular organisms\|d_Bacteria\|p_Proteobacteria\| |
| 0.369919 | 361 | S | 36809 | s_*Mycobacteroides abscessus* | 9 | -_root\|-_cellular organisms\|d_Bacteria\|-_Terrabacteria gr |
| 0.324832 | 317 | S | 556325 | s_*Neomicrococcus aestuarii* | 9 | -_root\|-_cellular organisms\|d_Bacteria\|-_Terrabacteria gr |
| 0.252078 | 246 | S | 68259 | s_*Streptomyces purpurogeneiscle* | 9 | -_root\|-_cellular organisms\|d_Bacteria\|-_Terrabacteria gr |
| 0.328931 | 321 | S | 1895717 | s_*Armatimonadetes bacterium* | 6 | -_root\|-_cellular organisms\|d_Bacteria\|-_Terrabacteria gr |
| 0.36582 | 357 | S | 1005039 | s_*Fimbriimonas ginsengisoli* | 9 | -_root\|-_cellular organisms\|d_Bacteria\|-_Terrabacteria gr |
| 0.294091 | 287 | S | 2052166 | s_*Candidatus Melainabacteria ba* | 7 | -_root\|-_cellular organisms\|d_Bacteria\|-_Terrabacteria gr |
| 0.462142 | 451 | S | 1004304 | s_*Hydrotalea sandarakina* | 10 | -_root\|-_cellular organisms\|d_Bacteria\|-_FCB group\|-_Ba |
