## supplementary data 3 for "Trophic Interactions of *Anopheles Gambiae* Mosquito Larvae in Aquatic Ecosystem: A Metagenomics Approach"

### *The family of eukaryotes found associated with the selected predators' diet*

| Predator | Rel. abundance | CladeReads | TaxRank | TaxID | Name | TaxLineage |
|---|---|---|---|---|---|---|
| Toxorynchithes | 80.3289 | 291619 | D | 2759 | d_Eukaryota | -_root\|-_cellular organisms\|d_Eukaryota |
| | 59.8844 | 217511 | F | 7157 | f_Culicidae | -_root\|-_cellular organisms\|d_Eukaryota\|-_Opisthokonta\|k_Metazoa\|-_Eumetazoa |
| | 0.00743355 | 27 | F | 7149 | f_Chironomidae | -_root\|-_cellular organisms\|d_Eukaryota\|-_Opisthokonta\|k_Metazoa\|-_Eumetazoa |
| | 0.0509336 | 185 | F | 7197 | f_Psychodidae | -_root\|-_cellular organisms\|d_Eukaryota\|-_Opisthokonta\|k_Metazoa\|-_Eumetazoa |
| | 2.107 | 7653 | F | 7214 | f_Drosophilidae | -_root\|-_cellular organisms\|d_Eukaryota\|-_Opisthokonta\|k_Metazoa\|-_Eumetazoa |
| | 0.0250538 | 91 | F | 7211 | f_Tephritidae | -_root\|-_cellular organisms\|d_Eukaryota\|-_Opisthokonta\|k_Metazoa\|-_Eumetazoa |
| | 0.0256045 | 93 | F | 7392 | f_Glossinidae | -_root\|-_cellular organisms\|d_Eukaryota\|-_Opisthokonta\|k_Metazoa\|-_Eumetazoa |
| | 0.0203734 | 74 | F | 7371 | f_Calliphoridae | -_root\|-_cellular organisms\|d_Eukaryota\|-_Opisthokonta\|k_Metazoa\|-_Eumetazoa |
| | 0.00853482 | 31 | F | 7366 | f_Muscidae | -_root\|-_cellular organisms\|d_Eukaryota\|-_Opisthokonta\|k_Metazoa\|-_Eumetazoa |
| | 0.00357912 | 13 | F | 36164 | f_Phoridae | -_root\|-_cellular organisms\|d_Eukaryota\|-_Opisthokonta\|k_Metazoa\|-_Eumetazoa |
| | 0.206488 | 750 | F | 36668 | f_Formicidae | -_root\|-_cellular organisms\|d_Eukaryota\|-_Opisthokonta\|k_Metazoa\|-_Eumetazoa |
| | 0.00633229 | 23 | F | 7458 | f_Apidae | -_root\|-_cellular organisms\|d_Eukaryota\|-_Opisthokonta\|k_Metazoa\|-_Eumetazoa |
| | 0.00357912 | 13 | F | 77572 | f_Halictidae | -_root\|-_cellular organisms\|d_Eukaryota\|-_Opisthokonta\|k_Metazoa\|-_Eumetazoa |
| | 0.00770887 | 28 | F | 7423 | f_Pteromalidae | -_root\|-_cellular organisms\|d_Eukaryota\|-_Opisthokonta\|k_Metazoa\|-_Eumetazoa |
| | 0.0300095 | 109 | F | 7049 | f_Lampyridae | -_root\|-_cellular organisms\|d_Eukaryota\|-_Opisthokonta\|k_Metazoa\|-_Eumetazoa |
| | 0.00770887 | 28 | F | 7065 | f_Tenebrionidae | -_root\|-_cellular organisms\|d_Eukaryota\|-_Opisthokonta\|k_Metazoa\|-_Eumetazoa |
| | 0.00770887 | 28 | F | 34667 | f_Cerambycidae | -_root\|-_cellular organisms\|d_Eukaryota\|-_Opisthokonta\|k_Metazoa\|-_Eumetazoa |
| | 0.0151424 | 55 | F | 7100 | f_Noctuidae | -_root\|-_cellular organisms\|d_Eukaryota\|-_Opisthokonta\|k_Metazoa\|-_Eumetazoa |
| | 0.00385443 | 14 | F | 7143 | f_Papilionidae | -_root\|-_cellular organisms\|d_Eukaryota\|-_Opisthokonta\|k_Metazoa\|-_Eumetazoa |
| | 0.00385443 | 14 | F | 82593 | f_Geometridae | -_root\|-_cellular organisms\|d_Eukaryota\|-_Opisthokonta\|k_Metazoa\|-_Eumetazoa |
| | 0.109025 | 396 | F | 27482 | f_Aphididae | -_root\|-_cellular organisms\|d_Eukaryota\|-_Opisthokonta\|k_Metazoa\|-_Eumetazoa |
| | 0.00633229 | 23 | F | 1585420 | f_Liviidae | -_root\|-_cellular organisms\|d_Eukaryota\|-_Opisthokonta\|k_Metazoa\|-_Eumetazoa |
| | 0.0189969 | 69 | F | 30083 | f_Miridae | -_root\|-_cellular organisms\|d_Eukaryota\|-_Opisthokonta\|k_Metazoa\|-_Eumetazoa |
| | 0.0140412 | 51 | F | 27479 | f_Reduviidae | -_root\|-_cellular organisms\|d_Eukaryota\|-_Opisthokonta\|k_Metazoa\|-_Eumetazoa |
| | 0.0145918 | 53 | F | 30102 | f_Cicadellidae | -_root\|-_cellular organisms\|d_Eukaryota\|-_Opisthokonta\|k_Metazoa\|-_Eumetazoa |
| | 0.011288 | 41 | F | 1049651 | f_Ectobiidae | -_root\|-_cellular organisms\|d_Eukaryota\|-_Opisthokonta\|k_Metazoa\|-_Eumetazoa |
| | 0.00357912 | 13 | F | 46562 | f_Kalotermitidae | -_root\|-_cellular organisms\|d_Eukaryota\|-_Opisthokonta\|k_Metazoa\|-_Eumetazoa |
| | 0.0143165 | 52 | F | 36141 | f_Isotomidae | -_root\|-_cellular organisms\|d_Eukaryota\|-_Opisthokonta\|k_Metazoa\|-_Eumetazoa |
| | 0.0203734 | 74 | F | 77658 | f_Daphniidae | -_root\|-_cellular organisms\|d_Eukaryota\|-_Opisthokonta\|k_Metazoa\|-_Eumetazoa |
| | 0.00798419 | 29 | F | 72034 | f_Caligidae | -_root\|-_cellular organisms\|d_Eukaryota\|-_Opisthokonta\|k_Metazoa\|-_Eumetazoa |
| | 0.0126646 | 46 | F | 6939 | f_Ixodidae | -_root\|-_cellular organisms\|d_Eukaryota\|-_Opisthokonta\|k_Metazoa\|-_Eumetazoa |
| | 0.00357912 | 13 | F | 6936 | f_Argasidae | -_root\|-_cellular organisms\|d_Eukaryota\|-_Opisthokonta\|k_Metazoa\|-_Eumetazoa |
| | 0.0101867 | 37 | F | 450948 | f_Nephilidae | -_root\|-_cellular organisms\|d_Eukaryota\|-_Opisthokonta\|k_Metazoa\|-_Eumetazoa |
| | 0.0049557 | 18 | F | 175333 | f_Eresidae | -_root\|-_cellular organisms\|d_Eukaryota\|-_Opisthokonta\|k_Metazoa\|-_Eumetazoa |
| | 0.320744 | 1165 | F | 6296 | f_Onchocercidae | -_root\|-_cellular organisms\|d_Eukaryota\|-_Opisthokonta\|k_Metazoa\|-_Eumetazoa |
| | 0.0154177 | 56 | F | 6243 | f_Rhabditidae | -_root\|-_cellular organisms\|d_Eukaryota\|-_Opisthokonta\|k_Metazoa\|-_Eumetazoa |
| | 0.00853482 | 31 | F | 6246 | f_Strongyloididae | -_root\|-_cellular organisms\|d_Eukaryota\|-_Opisthokonta\|k_Metazoa\|-_Eumetazoa |
| | 0.00578165 | 21 | F | 33278 | f_Ancylostomatic | -_root\|-_cellular organisms\|d_Eukaryota\|-_Opisthokonta\|k_Metazoa\|-_Eumetazoa |
| | 0.00770887 | 28 | F | 6332 | f_Trichinellidae | -_root\|-_cellular organisms\|d_Eukaryota\|-_Opisthokonta\|k_Metazoa\|-_Eumetazoa |
| | 0.00578165 | 21 | F | 119093 | f_Trichuridae | -_root\|-_cellular organisms\|d_Eukaryota\|-_Opisthokonta\|k_Metazoa\|-_Eumetazoa |
| | 0.00412975 | 15 | F | 31245 | f_Schistosomatid | -_root\|-_cellular organisms\|d_Eukaryota\|-_Opisthokonta\|k_Metazoa\|-_Eumetazoa |
| | 0.00412975 | 15 | F | 39216 | f_Macrostomidae | -_root\|-_cellular organisms\|d_Eukaryota\|-_Opisthokonta\|k_Metazoa\|-_Eumetazoa |
| | 0.00688292 | 25 | F | 54973 | f_Ampullariidae | -_root\|-_cellular organisms\|d_Eukaryota\|-_Opisthokonta\|k_Metazoa\|-_Eumetazoa |
| | 0.0033038 | 12 | F | 31181 | f_Strongylocentr | -_root\|-_cellular organisms\|d_Eukaryota\|-_Opisthokonta\|k_Metazoa\|-_Eumetazoa |
| | 0.00578165 | 21 | F | 46729 | f_Pocilloporidae | -_root\|-_cellular organisms\|d_Eukaryota\|-_Opisthokonta\|k_Metazoa\|-_Eumetazoa |
| | 0.0049557 | 18 | F | 6080 | f_Hydridae | -_root\|-_cellular organisms\|d_Eukaryota\|-_Opisthokonta\|k_Metazoa\|-_Eumetazoa |
| | 0.0033038 | 12 | F | 1352827 | f_Chytriomycetac | -_root\|-_cellular organisms\|d_Eukaryota\|-_Opisthokonta\|k_Fungi\|-_Fungi incertae |
| | 0.27807 | 1010 | F | 1131492 | f_Aspergillaceae | -_root\|-_cellular organisms\|d_Eukaryota\|-_Opisthokonta\|k_Fungi\|-_Dikarya\|p_Asc |
| | 0.0622216 | 226 | F | 34397 | f_Clavicipitaceae | -_root\|-_cellular organisms\|d_Eukaryota\|-_Opisthokonta\|k_Fungi\|-_Dikarya\|p_Asc |
| | 0.00302848 | 11 | F | 28556 | f_Pleosporaceae | -_root\|-_cellular organisms\|d_Eukaryota\|-_Opisthokonta\|k_Fungi\|-_Dikarya\|p_Asc |
| | 0.00302848 | 11 | F | 5250 | f_Ceratobasidiac | -_root\|-_cellular organisms\|d_Eukaryota\|-_Opisthokonta\|k_Fungi\|-_Dikarya\|p_Bas |

| Predator | Rel. abundance | CladeReads | TaxRank | TaxID | Name | TaxLineage |
|---|---|---|---|---|---|---|
| **Copepod** | 24.2404 | 15776 | D | 2759 | d_Eukaryota | -_root\|-_cellular organisms\|d_Eukaryota |
| | 0.150654 | 101 | F | 7157 | f_Culicidae | -_root\|-_cellular organisms\|d_Eukaryota\|-_Opisthokonta\|k_Metazoa\|-_Eumetazoa |
| | 0.00745812 | 5 | F | 7149 | f_Chironomidae | -_root\|-_cellular organisms\|d_Eukaryota\|-_Opisthokonta\|k_Metazoa\|-_Eumetazoa |
| | 0.0223744 | 15 | F | 7197 | f_Psychodidae | -_root\|-_cellular organisms\|d_Eukaryota\|-_Opisthokonta\|k_Metazoa\|-_Eumetazoa |
| | 0.0208827 | 14 | F | 7214 | f_Drosophilidae | -_root\|-_cellular organisms\|d_Eukaryota\|-_Opisthokonta\|k_Metazoa\|-_Eumetazoa |
| | 0.0880058 | 59 | F | 36668 | f_Formicidae | -_root\|-_cellular organisms\|d_Eukaryota\|-_Opisthokonta\|k_Metazoa\|-_Eumetazoa |
| | 0.0164079 | 11 | F | 77572 | f_Halictidae | -_root\|-_cellular organisms\|d_Eukaryota\|-_Opisthokonta\|k_Metazoa\|-_Eumetazoa |
| | 0.00894975 | 6 | F | 7458 | f_Apidae | -_root\|-_cellular organisms\|d_Eukaryota\|-_Opisthokonta\|k_Metazoa\|-_Eumetazoa |
| | 0.00894975 | 6 | F | 7423 | f_Pteromalidae | -_root\|-_cellular organisms\|d_Eukaryota\|-_Opisthokonta\|k_Metazoa\|-_Eumetazoa |
| | 0.0149162 | 10 | F | 7100 | f_Noctuidae | -_root\|-_cellular organisms\|d_Eukaryota\|-_Opisthokonta\|k_Metazoa\|-_Eumetazoa |
| | 0.00745812 | 5 | F | 7089 | f_Bombycidae | -_root\|-_cellular organisms\|d_Eukaryota\|-_Opisthokonta\|k_Metazoa\|-_Eumetazoa |
| | 0.011933 | 8 | F | 7065 | f_Tenebrionidae | -_root\|-_cellular organisms\|d_Eukaryota\|-_Opisthokonta\|k_Metazoa\|-_Eumetazoa |
| | 0.00894975 | 6 | F | 7049 | f_Lampyridae | -_root\|-_cellular organisms\|d_Eukaryota\|-_Opisthokonta\|k_Metazoa\|-_Eumetazoa |
| | 0.186453 | 125 | F | 30102 | f_Cicadellidae | -_root\|-_cellular organisms\|d_Eukaryota\|-_Opisthokonta\|k_Metazoa\|-_Eumetazoa |
| | 0.083531 | 56 | F | 27482 | f_Aphididae | -_root\|-_cellular organisms\|d_Eukaryota\|-_Opisthokonta\|k_Metazoa\|-_Eumetazoa |
| | 0.0536985 | 36 | F | 30083 | f_Miridae | -_root\|-_cellular organisms\|d_Eukaryota\|-_Opisthokonta\|k_Metazoa\|-_Eumetazoa |
| | 0.0134246 | 9 | F | 27479 | f_Reduviidae | -_root\|-_cellular organisms\|d_Eukaryota\|-_Opisthokonta\|k_Metazoa\|-_Eumetazoa |
| | 0.00447487 | 3 | F | 7533 | f_Lygaeidae | -_root\|-_cellular organisms\|d_Eukaryota\|-_Opisthokonta\|k_Metazoa\|-_Eumetazoa |
| | 0.0372906 | 25 | F | 46562 | f_Kalotermitidae | -_root\|-_cellular organisms\|d_Eukaryota\|-_Opisthokonta\|k_Metazoa\|-_Eumetazoa |
| | 0.0059665 | 4 | F | 1049651 | f_Ectobiidae | -_root\|-_cellular organisms\|d_Eukaryota\|-_Opisthokonta\|k_Metazoa\|-_Eumetazoa |
| | 0.00298325 | 2 | F | 48704 | f_Entomobryidae | -_root\|-_cellular organisms\|d_Eukaryota\|-_Opisthokonta\|k_Metazoa\|-_Eumetazoa |
| | 0.283409 | 190 | F | 72034 | f_Caligidae | -_root\|-_cellular organisms\|d_Eukaryota\|-_Opisthokonta\|k_Metazoa\|-_Eumetazoa |
| | 0.00745812 | 5 | F | 512265 | f_Harpacticidae | -_root\|-_cellular organisms\|d_Eukaryota\|-_Opisthokonta\|k_Metazoa\|-_Eumetazoa |
| | 0.0059665 | 4 | F | 564998 | f_Cyclopettidae | -_root\|-_cellular organisms\|d_Eukaryota\|-_Opisthokonta\|k_Metazoa\|-_Eumetazoa |
| | 0.0149162 | 10 | F | 121147 | f_Acartiidae | -_root\|-_cellular organisms\|d_Eukaryota\|-_Opisthokonta\|k_Metazoa\|-_Eumetazoa |
| | 0.011933 | 8 | F | 163474 | f_Pseudodiapton | -_root\|-_cellular organisms\|d_Eukaryota\|-_Opisthokonta\|k_Metazoa\|-_Eumetazoa |
| | 0.00447487 | 3 | F | 163477 | f_Tortanidae | -_root\|-_cellular organisms\|d_Eukaryota\|-_Opisthokonta\|k_Metazoa\|-_Eumetazoa |
| | 0.0387822 | 26 | F | 6757 | f_Portunidae | -_root\|-_cellular organisms\|d_Eukaryota\|-_Opisthokonta\|k_Metazoa\|-_Eumetazoa |
| | 0.0387822 | 26 | G | 6760 | g_Scylla | -_root\|-_cellular organisms\|d_Eukaryota\|-_Opisthokonta\|k_Metazoa\|-_Eumetazoa |
| | 0.0387822 | 26 | S | 85551 | s_Scylla olivacea | -_root\|-_cellular organisms\|d_Eukaryota\|-_Opisthokonta\|k_Metazoa\|-_Eumetazoa |
| | 0.0059665 | 4 | F | 134545 | f_Lysianassidae | -_root\|-_cellular organisms\|d_Eukaryota\|-_Opisthokonta\|k_Metazoa\|-_Eumetazoa |
| | 0.0536985 | 36 | F | 77658 | f_Daphniidae | -_root\|-_cellular organisms\|d_Eukaryota\|-_Opisthokonta\|k_Metazoa\|-_Eumetazoa |
| | 0.00447487 | 3 | F | 126955 | f_Linotaeniidae | -_root\|-_cellular organisms\|d_Eukaryota\|-_Opisthokonta\|k_Metazoa\|-_Eumetazoa |
| | 0.108889 | 73 | F | 6939 | f_Ixodidae | -_root\|-_cellular organisms\|d_Eukaryota\|-_Opisthokonta\|k_Metazoa\|-_Eumetazoa |
| | 0.0611566 | 41 | F | 6936 | f_Argasidae | -_root\|-_cellular organisms\|d_Eukaryota\|-_Opisthokonta\|k_Metazoa\|-_Eumetazoa |
| | 0.023866 | 16 | F | 450948 | f_Nephilidae | -_root\|-_cellular organisms\|d_Eukaryota\|-_Opisthokonta\|k_Metazoa\|-_Eumetazoa |
| | 0.00447487 | 3 | F | 34643 | f_Theridiidae | -_root\|-_cellular organisms\|d_Eukaryota\|-_Opisthokonta\|k_Metazoa\|-_Eumetazoa |
| | 0.00447487 | 3 | F | 175333 | f_Eresidae | -_root\|-_cellular organisms\|d_Eukaryota\|-_Opisthokonta\|k_Metazoa\|-_Eumetazoa |
| | 0.047732 | 32 | F | 6246 | f_Strongyloidida | -_root\|-_cellular organisms\|d_Eukaryota\|-_Opisthokonta\|k_Metazoa\|-_Eumetazoa |
| | 0.0134246 | 9 | F | 33253 | f_Steinernematic | -_root\|-_cellular organisms\|d_Eukaryota\|-_Opisthokonta\|k_Metazoa\|-_Eumetazoa |
| | 0.0492236 | 33 | F | 6243 | f_Rhabditidae | -_root\|-_cellular organisms\|d_Eukaryota\|-_Opisthokonta\|k_Metazoa\|-_Eumetazoa |
| | 0.0417655 | 28 | F | 119093 | f_Trichuridae | -_root\|-_cellular organisms\|d_Eukaryota\|-_Opisthokonta\|k_Metazoa\|-_Eumetazoa |
| | 0.0208827 | 14 | F | 6332 | f_Trichinellidae | -_root\|-_cellular organisms\|d_Eukaryota\|-_Opisthokonta\|k_Metazoa\|-_Eumetazoa |
| | 0.0850226 | 57 | F | 39216 | f_Macrostomidae | -_root\|-_cellular organisms\|d_Eukaryota\|-_Opisthokonta\|k_Metazoa\|-_Eumetazoa |
| | 0.0149162 | 10 | F | 99737 | f_Echinostomatic | -_root\|-_cellular organisms\|d_Eukaryota\|-_Opisthokonta\|k_Metazoa\|-_Eumetazoa |
| | 0.0104414 | 7 | F | 31245 | f_Schistosomatid | -_root\|-_cellular organisms\|d_Eukaryota\|-_Opisthokonta\|k_Metazoa\|-_Eumetazoa |
| | 0.0104414 | 7 | F | 6196 | f_Opisthorchiida | -_root\|-_cellular organisms\|d_Eukaryota\|-_Opisthokonta\|k_Metazoa\|-_Eumetazoa |
| | 0.0178995 | 12 | F | 6214 | f_Hymenolepidic | -_root\|-_cellular organisms\|d_Eukaryota\|-_Opisthokonta\|k_Metazoa\|-_Eumetazoa |
| | 0.0432571 | 29 | F | 51292 | f_Capitellidae | -_root\|-_cellular organisms\|d_Eukaryota\|-_Opisthokonta\|k_Metazoa\|-_Eumetazoa |
| | 0.011933 | 8 | F | 6407 | f_Glossiphoniida | -_root\|-_cellular organisms\|d_Eukaryota\|-_Opisthokonta\|k_Metazoa\|-_Eumetazoa |
| | 0.0223744 | 15 | F | 6647 | f_Octopodidae | -_root\|-_cellular organisms\|d_Eukaryota\|-_Opisthokonta\|k_Metazoa\|-_Eumetazoa |
| | 0.00745812 | 5 | F | 33491 | f_Lingulidae | -_root\|-_cellular organisms\|d_Eukaryota\|-_Opisthokonta\|k_Metazoa\|-_Eumetazoa |
| | 0.105905 | 71 | F | 7687 | f_Stichopodidae | -_root\|-_cellular organisms\|d_Eukaryota\|-_Opisthokonta\|k_Metazoa\|-_Eumetazoa |
| | 0.0462404 | 31 | F | 31181 | f_Strongylocentr | -_root\|-_cellular organisms\|d_Eukaryota\|-_Opisthokonta\|k_Metazoa\|-_Eumetazoa |
| | 0.0447487 | 30 | F | 6080 | f_Hydridae | -_root\|-_cellular organisms\|d_Eukaryota\|-_Opisthokonta\|k_Metazoa\|-_Eumetazoa |
| | 0.0149162 | 10 | F | 46729 | f_Pocilloporidae | -_root\|-_cellular organisms\|d_Eukaryota\|-_Opisthokonta\|k_Metazoa\|-_Eumetazoa |
| | 0.0134246 | 9 | F | 45349 | f_Edwardsiidae | -_root\|-_cellular organisms\|d_Eukaryota\|-_Opisthokonta\|k_Metazoa\|-_Eumetazoa |
| | 0.0268492 | 18 | F | 35587 | f_Myxobolidae | -_root\|-_cellular organisms\|d_Eukaryota\|-_Opisthokonta\|k_Metazoa\|-_Eumetazoa |
| | 0.0253576 | 17 | F | 178475 | f_Niphatidae | -_root\|-_cellular organisms\|d_Eukaryota\|-_Opisthokonta\|k_Metazoa\|p_Porifera\|c_ |
| | 0.0134246 | 9 | F | 34397 | f_Clavicipitaceae | -_root\|-_cellular organisms\|d_Eukaryota\|-_Opisthokonta\|k_Fungi\|-_Dikarya\|p_Asc |
| | 0.0178995 | 12 | F | 28556 | f_Pleosporaceae | -_root\|-_cellular organisms\|d_Eukaryota\|-_Opisthokonta\|k_Fungi\|-_Dikarya\|p_Asc |
| | 0.00745812 | 5 | F | 1131492 | f_Aspergillaceae | -_root\|-_cellular organisms\|d_Eukaryota\|-_Opisthokonta\|k_Fungi\|-_Dikarya\|p_Asc |

<table>
<tr><th>Predator</th><th>Rel. abundance</th><th>CladeReads</th><th>TaxRank</th><th>TaxID</th><th>Name</th><th>TaxLineage</th></tr>
<tr><td>**Backswimmer**</td><td>24.0468</td><td>47215</td><td>D</td><td>2759</td><td>d_Eukaryota</td><td>-_root|-_cellular organisms|d_Eukaryota</td></tr>
<tr><td></td><td>0.175035</td><td>350</td><td>F</td><td>7157</td><td>f_Culicidae</td><td>-_root|-_cellular organisms|d_Eukaryota|-_Opisthokonta|k_Metazoa|-_Eumetazoa</td></tr>
<tr><td></td><td>0.0095019</td><td>19</td><td>F</td><td>41811</td><td>f_Chaoboridae</td><td>-_root|-_cellular organisms|d_Eukaryota|-_Opisthokonta|k_Metazoa|-_Eumetazoa</td></tr>
<tr><td></td><td>0.0160032</td><td>32</td><td>F</td><td>7149</td><td>f_Chironomidae</td><td>-_root|-_cellular organisms|d_Eukaryota|-_Opisthokonta|k_Metazoa|-_Eumetazoa</td></tr>
<tr><td></td><td>0.0580116</td><td>116</td><td>F</td><td>7197</td><td>f_Psychodidae</td><td>-_root|-_cellular organisms|d_Eukaryota|-_Opisthokonta|k_Metazoa|-_Eumetazoa</td></tr>
<tr><td></td><td>0.0060012</td><td>12</td><td>F</td><td>33406</td><td>f_Cecidomyiidae</td><td>-_root|-_cellular organisms|d_Eukaryota|-_Opisthokonta|k_Metazoa|-_Eumetazoa</td></tr>
<tr><td></td><td>0.0310062</td><td>62</td><td>F</td><td>7214</td><td>f_Drosophilidae</td><td>-_root|-_cellular organisms|d_Eukaryota|-_Opisthokonta|k_Metazoa|-_Eumetazoa</td></tr>
<tr><td></td><td>0.0220044</td><td>44</td><td>F</td><td>7211</td><td>f_Tephritidae</td><td>-_root|-_cellular organisms|d_Eukaryota|-_Opisthokonta|k_Metazoa|-_Eumetazoa</td></tr>
<tr><td></td><td>0.0130026</td><td>26</td><td>F</td><td>7392</td><td>f_Glossinidae</td><td>-_root|-_cellular organisms|d_Eukaryota|-_Opisthokonta|k_Metazoa|-_Eumetazoa</td></tr>
<tr><td></td><td>0.0075015</td><td>15</td><td>F</td><td>7366</td><td>f_Muscidae</td><td>-_root|-_cellular organisms|d_Eukaryota|-_Opisthokonta|k_Metazoa|-_Eumetazoa</td></tr>
<tr><td></td><td>0.0055011</td><td>11</td><td>F</td><td>7205</td><td>f_Tabanidae</td><td>-_root|-_cellular organisms|d_Eukaryota|-_Opisthokonta|k_Metazoa|-_Eumetazoa</td></tr>
<tr><td></td><td>0.162533</td><td>325</td><td>F</td><td>36668</td><td>f_Formicidae</td><td>-_root|-_cellular organisms|d_Eukaryota|-_Opisthokonta|k_Metazoa|-_Eumetazoa</td></tr>
<tr><td></td><td>0.0215043</td><td>43</td><td>F</td><td>7458</td><td>f_Apidae</td><td>-_root|-_cellular organisms|d_Eukaryota|-_Opisthokonta|k_Metazoa|-_Eumetazoa</td></tr>
<tr><td></td><td>0.0065013</td><td>13</td><td>F</td><td>77572</td><td>f_Halictidae</td><td>-_root|-_cellular organisms|d_Eukaryota|-_Opisthokonta|k_Metazoa|-_Eumetazoa</td></tr>
<tr><td></td><td>0.0335067</td><td>67</td><td>F</td><td>7423</td><td>f_Pteromalidae</td><td>-_root|-_cellular organisms|d_Eukaryota|-_Opisthokonta|k_Metazoa|-_Eumetazoa</td></tr>
<tr><td></td><td>0.0095019</td><td>19</td><td>F</td><td>7402</td><td>f_Braconidae</td><td>-_root|-_cellular organisms|d_Eukaryota|-_Opisthokonta|k_Metazoa|-_Eumetazoa</td></tr>
<tr><td></td><td>0.0940188</td><td>188</td><td>F</td><td>7049</td><td>f_Lampyridae</td><td>-_root|-_cellular organisms|d_Eukaryota|-_Opisthokonta|k_Metazoa|-_Eumetazoa</td></tr>
<tr><td></td><td>0.0255051</td><td>51</td><td>F</td><td>50527</td><td>f_Buprestidae</td><td>-_root|-_cellular organisms|d_Eukaryota|-_Opisthokonta|k_Metazoa|-_Eumetazoa</td></tr>
<tr><td></td><td>0.055011</td><td>110</td><td>F</td><td>7065</td><td>f_Tenebrionidae</td><td>-_root|-_cellular organisms|d_Eukaryota|-_Opisthokonta|k_Metazoa|-_Eumetazoa</td></tr>
<tr><td></td><td>0.0190038</td><td>38</td><td>F</td><td>7042</td><td>f_Curculionidae</td><td>-_root|-_cellular organisms|d_Eukaryota|-_Opisthokonta|k_Metazoa|-_Eumetazoa</td></tr>
<tr><td></td><td>0.0130026</td><td>26</td><td>F</td><td>34667</td><td>f_Cerambycidae</td><td>-_root|-_cellular organisms|d_Eukaryota|-_Opisthokonta|k_Metazoa|-_Eumetazoa</td></tr>
<tr><td></td><td>0.0125025</td><td>25</td><td>F</td><td>7055</td><td>f_Scarabaeidae</td><td>-_root|-_cellular organisms|d_Eukaryota|-_Opisthokonta|k_Metazoa|-_Eumetazoa</td></tr>
<tr><td></td><td>0.0575115</td><td>115</td><td>F</td><td>7100</td><td>f_Noctuidae</td><td>-_root|-_cellular organisms|d_Eukaryota|-_Opisthokonta|k_Metazoa|-_Eumetazoa</td></tr>
<tr><td></td><td>0.0125025</td><td>25</td><td>F</td><td>7143</td><td>f_Papilionidae</td><td>-_root|-_cellular organisms|d_Eukaryota|-_Opisthokonta|k_Metazoa|-_Eumetazoa</td></tr>
<tr><td></td><td>0.0085017</td><td>17</td><td>F</td><td>33415</td><td>f_Nymphalidae</td><td>-_root|-_cellular organisms|d_Eukaryota|-_Opisthokonta|k_Metazoa|-_Eumetazoa</td></tr>
<tr><td></td><td>0.010002</td><td>20</td><td>F</td><td>7089</td><td>f_Bombycidae</td><td>-_root|-_cellular organisms|d_Eukaryota|-_Opisthokonta|k_Metazoa|-_Eumetazoa</td></tr>
<tr><td></td><td>0.010002</td><td>20</td><td>F</td><td>82593</td><td>f_Geometridae</td><td>-_root|-_cellular organisms|d_Eukaryota|-_Opisthokonta|k_Metazoa|-_Eumetazoa</td></tr>
<tr><td></td><td>0.380576</td><td>761</td><td>F</td><td>27482</td><td>f_Aphididae</td><td>-_root|-_cellular organisms|d_Eukaryota|-_Opisthokonta|k_Metazoa|-_Eumetazoa</td></tr>
<tr><td></td><td>0.0740148</td><td>148</td><td>F</td><td>1585420</td><td>f_Liviidae</td><td>-_root|-_cellular organisms|d_Eukaryota|-_Opisthokonta|k_Metazoa|-_Eumetazoa</td></tr>
<tr><td></td><td>0.291558</td><td>583</td><td>F</td><td>30102</td><td>f_Cicadellidae</td><td>-_root|-_cellular organisms|d_Eukaryota|-_Opisthokonta|k_Metazoa|-_Eumetazoa</td></tr>
<tr><td></td><td>0.0560112</td><td>112</td><td>F</td><td>139606</td><td>f_Clastopteridae</td><td>-_root|-_cellular organisms|d_Eukaryota|-_Opisthokonta|k_Metazoa|-_Eumetazoa</td></tr>
<tr><td></td><td>0.0055011</td><td>11</td><td>F</td><td>33362</td><td>f_Delphacidae</td><td>-_root|-_cellular organisms|d_Eukaryota|-_Opisthokonta|k_Metazoa|-_Eumetazoa</td></tr>
<tr><td></td><td>0.126025</td><td>252</td><td>F</td><td>27479</td><td>f_Reduviidae</td><td>-_root|-_cellular organisms|d_Eukaryota|-_Opisthokonta|k_Metazoa|-_Eumetazoa</td></tr>
<tr><td></td><td>0.106021</td><td>212</td><td>F</td><td>30083</td><td>f_Miridae</td><td>-_root|-_cellular organisms|d_Eukaryota|-_Opisthokonta|k_Metazoa|-_Eumetazoa</td></tr>
<tr><td></td><td>0.0330066</td><td>66</td><td>F</td><td>7533</td><td>f_Lygaeidae</td><td>-_root|-_cellular organisms|d_Eukaryota|-_Opisthokonta|k_Metazoa|-_Eumetazoa</td></tr>
<tr><td></td><td>0.0720144</td><td>144</td><td>F</td><td>121221</td><td>f_Pediculidae</td><td>-_root|-_cellular organisms|d_Eukaryota|-_Opisthokonta|k_Metazoa|-_Eumetazoa</td></tr>
<tr><td></td><td>0.241048</td><td>482</td><td>F</td><td>46562</td><td>f_Kalotermitidae</td><td>-_root|-_cellular organisms|d_Eukaryota|-_Opisthokonta|k_Metazoa|-_Eumetazoa</td></tr>
<tr><td></td><td>0.0775155</td><td>155</td><td>F</td><td>7501</td><td>f_Termopsidae</td><td>-_root|-_cellular organisms|d_Eukaryota|-_Opisthokonta|k_Metazoa|-_Eumetazoa</td></tr>
<tr><td></td><td>0.0035007</td><td>7</td><td>F</td><td>36985</td><td>f_Rhinotermitidae</td><td>-_root|-_cellular organisms|d_Eukaryota|-_Opisthokonta|k_Metazoa|-_Eumetazoa</td></tr>
<tr><td></td><td>0.168034</td><td>336</td><td>F</td><td>1049651</td><td>f_Ectobiidae</td><td>-_root|-_cellular organisms|d_Eukaryota|-_Opisthokonta|k_Metazoa|-_Eumetazoa</td></tr>
<tr><td></td><td>0.109022</td><td>218</td><td>F</td><td>7002</td><td>f_Acrididae</td><td>-_root|-_cellular organisms|d_Eukaryota|-_Opisthokonta|k_Metazoa|-_Eumetazoa</td></tr>
<tr><td></td><td>0.0280056</td><td>56</td><td>F</td><td>172515</td><td>f_Baetidae</td><td>-_root|-_cellular organisms|d_Eukaryota|-_Opisthokonta|k_Metazoa|-_Eumetazoa</td></tr>
<tr><td></td><td>0.0035007</td><td>7</td><td>F</td><td>50581</td><td>f_Ephemeridae</td><td>-_root|-_cellular organisms|d_Eukaryota|-_Opisthokonta|k_Metazoa|-_Eumetazoa</td></tr>
<tr><td></td><td>0.0240048</td><td>48</td><td>F</td><td>48704</td><td>f_Entomobryidae</td><td>-_root|-_cellular organisms|d_Eukaryota|-_Opisthokonta|k_Metazoa|-_Eumetazoa</td></tr>
<tr><td></td><td>0.0225045</td><td>45</td><td>F</td><td>36141</td><td>f_Isotomidae</td><td>-_root|-_cellular organisms|d_Eukaryota|-_Opisthokonta|k_Metazoa|-_Eumetazoa</td></tr>
<tr><td></td><td>0.118024</td><td>236</td><td>F</td><td>77658</td><td>f_Daphniidae</td><td>-_root|-_cellular organisms|d_Eukaryota|-_Opisthokonta|k_Metazoa|-_Eumetazoa</td></tr>
<tr><td></td><td>0.0285057</td><td>57</td><td>F</td><td>6757</td><td>f_Portunidae</td><td>-_root|-_cellular organisms|d_Eukaryota|-_Opisthokonta|k_Metazoa|-_Eumetazoa</td></tr>
<tr><td></td><td>0.0070014</td><td>14</td><td>F</td><td>134545</td><td>f_Lysianassidae</td><td>-_root|-_cellular organisms|d_Eukaryota|-_Opisthokonta|k_Metazoa|-_Eumetazoa</td></tr>
<tr><td></td><td>0.0110022</td><td>22</td><td>F</td><td>72034</td><td>f_Caligidae</td><td>-_root|-_cellular organisms|d_Eukaryota|-_Opisthokonta|k_Metazoa|-_Eumetazoa</td></tr>
<tr><td></td><td>0.025005</td><td>50</td><td>F</td><td>126955</td><td>f_Linotaeniidae</td><td>-_root|-_cellular organisms|d_Eukaryota|-_Opisthokonta|k_Metazoa|-_Eumetazoa</td></tr>
<tr><td></td><td>0.127526</td><td>255</td><td>F</td><td>6939</td><td>f_Ixodidae</td><td>-_root|-_cellular organisms|d_Eukaryota|-_Opisthokonta|k_Metazoa|-_Eumetazoa</td></tr>
<tr><td></td><td>0.0160032</td><td>32</td><td>F</td><td>6936</td><td>f_Argasidae</td><td>-_root|-_cellular organisms|d_Eukaryota|-_Opisthokonta|k_Metazoa|-_Eumetazoa</td></tr>
<tr><td></td><td>0.0570114</td><td>114</td><td>F</td><td>450948</td><td>f_Nephilidae</td><td>-_root|-_cellular organisms|d_Eukaryota|-_Opisthokonta|k_Metazoa|-_Eumetazoa</td></tr>
<tr><td></td><td>0.0060012</td><td>12</td><td>F</td><td>34643</td><td>f_Theridiidae</td><td>-_root|-_cellular organisms|d_Eukaryota|-_Opisthokonta|k_Metazoa|-_Eumetazoa</td></tr>
<tr><td></td><td>0.0335067</td><td>67</td><td>F</td><td>175333</td><td>f_Eresidae</td><td>-_root|-_cellular organisms|d_Eukaryota|-_Opisthokonta|k_Metazoa|-_Eumetazoa</td></tr>
<tr><td></td><td>0.214043</td><td>428</td><td>F</td><td>6246</td><td>f_Strongyloididae</td><td>-_root|-_cellular organisms|d_Eukaryota|-_Opisthokonta|k_Metazoa|-_Eumetazoa</td></tr>
<tr><td></td><td>0.0315063</td><td>63</td><td>F</td><td>33253</td><td>f_Steinernematida</td><td>-_root|-_cellular organisms|d_Eukaryota|-_Opisthokonta|k_Metazoa|-_Eumetazoa</td></tr>
<tr><td></td><td>0.0815163</td><td>163</td><td>F</td><td>6243</td><td>f_Rhabditidae</td><td>-_root|-_cellular organisms|d_Eukaryota|-_Opisthokonta|k_Metazoa|-_Eumetazoa</td></tr>
<tr><td></td><td>0.0070014</td><td>14</td><td>F</td><td>6296</td><td>f_Onchocercidae</td><td>-_root|-_cellular organisms|d_Eukaryota|-_Opisthokonta|k_Metazoa|-_Eumetazoa</td></tr>
<tr><td></td><td>0.0195039</td><td>39</td><td>F</td><td>119093</td><td>f_Trichuridae</td><td>-_root|-_cellular organisms|d_Eukaryota|-_Opisthokonta|k_Metazoa|-_Eumetazoa</td></tr>
<tr><td></td><td>0.010002</td><td>20</td><td>F</td><td>6332</td><td>f_Trichinellidae</td><td>-_root|-_cellular organisms|d_Eukaryota|-_Opisthokonta|k_Metazoa|-_Eumetazoa</td></tr>
<tr><td></td><td>0.0165033</td><td>33</td><td>F</td><td>54973</td><td>f_Ampullariidae</td><td>-_root|-_cellular organisms|d_Eukaryota|-_Opisthokonta|k_Metazoa|-_Eumetazoa</td></tr>
<tr><td></td><td>0.005001</td><td>10</td><td>F</td><td>6524</td><td>f_Planorbidae</td><td>-_root|-_cellular organisms|d_Eukaryota|-_Opisthokonta|k_Metazoa|-_Eumetazoa</td></tr>
<tr><td></td><td>0.0035007</td><td>7</td><td>F</td><td>6540</td><td>f_Arionidae</td><td>-_root|-_cellular organisms|d_Eukaryota|-_Opisthokonta|k_Metazoa|-_Eumetazoa</td></tr>
<tr><td></td><td>0.0070014</td><td>14</td><td>F</td><td>69676</td><td>f_Lottiidae</td><td>-_root|-_cellular organisms|d_Eukaryota|-_Opisthokonta|k_Metazoa|-_Eumetazoa</td></tr>
<tr><td></td><td>0.0045009</td><td>9</td><td>F</td><td>6566</td><td>f_Pectinidae</td><td>-_root|-_cellular organisms|d_Eukaryota|-_Opisthokonta|k_Metazoa|-_Eumetazoa</td></tr>
<tr><td></td><td>0.0055011</td><td>11</td><td>F</td><td>39216</td><td>f_Macrostomidae</td><td>-_root|-_cellular organisms|d_Eukaryota|-_Opisthokonta|k_Metazoa|-_Eumetazoa</td></tr>
<tr><td></td><td>0.0080016</td><td>16</td><td>F</td><td>51292</td><td>f_Capitellidae</td><td>-_root|-_cellular organisms|d_Eukaryota|-_Opisthokonta|k_Metazoa|-_Eumetazoa</td></tr>
<tr><td></td><td>0.0075015</td><td>15</td><td>F</td><td>6407</td><td>f_Glossiphoniidae</td><td>-_root|-_cellular organisms|d_Eukaryota|-_Opisthokonta|k_Metazoa|-_Eumetazoa</td></tr>
<tr><td></td><td>0.0080016</td><td>16</td><td>F</td><td>33491</td><td>f_Lingulidae</td><td>-_root|-_cellular organisms|d_Eukaryota|-_Opisthokonta|k_Metazoa|-_Eumetazoa</td></tr>
<tr><td></td><td>0.035007</td><td>70</td><td>F</td><td>31181</td><td>f_Strongylocentro</td><td>-_root|-_cellular organisms|d_Eukaryota|-_Opisthokonta|k_Metazoa|-_Eumetazoa</td></tr>
<tr><td></td><td>0.0155031</td><td>31</td><td>F</td><td>7687</td><td>f_Stichopodidae</td><td>-_root|-_cellular organisms|d_Eukaryota|-_Opisthokonta|k_Metazoa|-_Eumetazoa</td></tr>
<tr><td></td><td>0.0215043</td><td>43</td><td>F</td><td>46729</td><td>f_Pocilloporidae</td><td>-_root|-_cellular organisms|d_Eukaryota|-_Opisthokonta|k_Metazoa|-_Eumetazoa</td></tr>
<tr><td></td><td>0.0080016</td><td>16</td><td>F</td><td>474943</td><td>f_Cordycipitaceae</td><td>-_root|-_cellular organisms|d_Eukaryota|-_Opisthokonta|k_Fungi|-_Dikarya|p_Asc</td></tr>
<tr><td></td><td>0.0135027</td><td>27</td><td>F</td><td>1131492</td><td>f_Aspergillaceae</td><td>-_root|-_cellular organisms|d_Eukaryota|-_Opisthokonta|k_Fungi|-_Dikarya|p_Asc</td></tr>
</table>

| Predator | Rel. abundance | CladeReads | TaxRank | TaxID | Name | TaxLineage |
|---|---|---|---|---|---|---|
| **Dragonfly** | 81.4466 | 54887 | D | 2759 | d_Eukaryota | -_root\|-_cellular organisms\|d_Eukaryota |
| | 1.35425 | 927 | F | 36668 | f_Formicidae | -_root\|-_cellular organisms\|d_Eukaryota\|-_Opisthokonta\|k_Metazoa\|-_Eumetazoa |
| | 0.327241 | 224 | F | 7458 | f_Apidae | -_root\|-_cellular organisms\|d_Eukaryota\|-_Opisthokonta\|k_Metazoa\|-_Eumetazoa |
| | 0.0613578 | 42 | F | 77572 | f_Halictidae | -_root\|-_cellular organisms\|d_Eukaryota\|-_Opisthokonta\|k_Metazoa\|-_Eumetazoa |
| | 0.214752 | 147 | F | 7423 | f_Pteromalidae | -_root\|-_cellular organisms\|d_Eukaryota\|-_Opisthokonta\|k_Metazoa\|-_Eumetazoa |
| | 0.0569751 | 39 | F | 7402 | f_Braconidae | -_root\|-_cellular organisms\|d_Eukaryota\|-_Opisthokonta\|k_Metazoa\|-_Eumetazoa |
| | 1.39662 | 956 | F | 7157 | f_Culicidae | -_root\|-_cellular organisms\|d_Eukaryota\|-_Opisthokonta\|k_Metazoa\|-_Eumetazoa |
| | 0.0394443 | 27 | F | 41811 | f_Chaoboridae | -_root\|-_cellular organisms\|d_Eukaryota\|-_Opisthokonta\|k_Metazoa\|-_Eumetazoa |
| | 0.0701232 | 48 | F | 7149 | f_Chironomidae | -_root\|-_cellular organisms\|d_Eukaryota\|-_Opisthokonta\|k_Metazoa\|-_Eumetazoa |
| | 0.122716 | 84 | F | 7197 | f_Psychodidae | -_root\|-_cellular organisms\|d_Eukaryota\|-_Opisthokonta\|k_Metazoa\|-_Eumetazoa |
| | 0.230822 | 158 | F | 7392 | f_Glossinidae | -_root\|-_cellular organisms\|d_Eukaryota\|-_Opisthokonta\|k_Metazoa\|-_Eumetazoa |
| | 0.118333 | 81 | F | 7371 | f_Calliphoridae | -_root\|-_cellular organisms\|d_Eukaryota\|-_Opisthokonta\|k_Metazoa\|-_Eumetazoa |
| | 0.0233744 | 16 | F | 7366 | f_Muscidae | -_root\|-_cellular organisms\|d_Eukaryota\|-_Opisthokonta\|k_Metazoa\|-_Eumetazoa |
| | 0.119794 | 82 | F | 7211 | f_Tephritidae | -_root\|-_cellular organisms\|d_Eukaryota\|-_Opisthokonta\|k_Metazoa\|-_Eumetazoa |
| | 0.112489 | 77 | F | 7214 | f_Drosophilidae | -_root\|-_cellular organisms\|d_Eukaryota\|-_Opisthokonta\|k_Metazoa\|-_Eumetazoa |
| | 0.0189917 | 13 | F | 36164 | f_Phoridae | -_root\|-_cellular organisms\|d_Eukaryota\|-_Opisthokonta\|k_Metazoa\|-_Eumetazoa |
| | 0.014609 | 10 | F | 7205 | f_Tabanidae | -_root\|-_cellular organisms\|d_Eukaryota\|-_Opisthokonta\|k_Metazoa\|-_Eumetazoa |
| | 0.255657 | 175 | F | 7065 | f_Tenebrionidae | -_root\|-_cellular organisms\|d_Eukaryota\|-_Opisthokonta\|k_Metazoa\|-_Eumetazoa |
| | 0.127098 | 87 | F | 7042 | f_Curculionidae | -_root\|-_cellular organisms\|d_Eukaryota\|-_Opisthokonta\|k_Metazoa\|-_Eumetazoa |
| | 0.058436 | 40 | F | 34667 | f_Cerambycidae | -_root\|-_cellular organisms\|d_Eukaryota\|-_Opisthokonta\|k_Metazoa\|-_Eumetazoa |
| | 0.30825 | 211 | F | 7049 | f_Lampyridae | -_root\|-_cellular organisms\|d_Eukaryota\|-_Opisthokonta\|k_Metazoa\|-_Eumetazoa |
| | 0.138785 | 95 | F | 50527 | f_Buprestidae | -_root\|-_cellular organisms\|d_Eukaryota\|-_Opisthokonta\|k_Metazoa\|-_Eumetazoa |
| | 0.073045 | 50 | F | 7055 | f_Scarabaeidae | -_root\|-_cellular organisms\|d_Eukaryota\|-_Opisthokonta\|k_Metazoa\|-_Eumetazoa |
| | 0.166542 | 114 | F | 7100 | f_Noctuidae | -_root\|-_cellular organisms\|d_Eukaryota\|-_Opisthokonta\|k_Metazoa\|-_Eumetazoa |
| | 0.0613578 | 42 | F | 7143 | f_Papilionidae | -_root\|-_cellular organisms\|d_Eukaryota\|-_Opisthokonta\|k_Metazoa\|-_Eumetazoa |
| | 0.0423661 | 29 | F | 33415 | f_Nymphalidae | -_root\|-_cellular organisms\|d_Eukaryota\|-_Opisthokonta\|k_Metazoa\|-_Eumetazoa |
| | 0.0423661 | 29 | F | 7089 | f_Bombycidae | -_root\|-_cellular organisms\|d_Eukaryota\|-_Opisthokonta\|k_Metazoa\|-_Eumetazoa |
| | 0.0248353 | 17 | F | 82593 | f_Geometridae | -_root\|-_cellular organisms\|d_Eukaryota\|-_Opisthokonta\|k_Metazoa\|-_Eumetazoa |
| | 0.0189917 | 13 | F | 32434 | f_Gelechiidae | -_root\|-_cellular organisms\|d_Eukaryota\|-_Opisthokonta\|k_Metazoa\|-_Eumetazoa |
| | 1.70925 | 1170 | F | 46562 | f_Kalotermitidae | -_root\|-_cellular organisms\|d_Eukaryota\|-_Opisthokonta\|k_Metazoa\|-_Eumetazoa |
| | 0.663248 | 454 | F | 7501 | f_Termopsidae | -_root\|-_cellular organisms\|d_Eukaryota\|-_Opisthokonta\|k_Metazoa\|-_Eumetazoa |
| | 0.0175308 | 12 | F | 36985 | f_Rhinotermitidae | -_root\|-_cellular organisms\|d_Eukaryota\|-_Opisthokonta\|k_Metazoa\|-_Eumetazoa |
| | 1.1658 | 798 | F | 1049651 | f_Ectobiidae | -_root\|-_cellular organisms\|d_Eukaryota\|-_Opisthokonta\|k_Metazoa\|-_Eumetazoa |
| | 0.0788885 | 54 | F | 7002 | f_Acrididae | -_root\|-_cellular organisms\|d_Eukaryota\|-_Opisthokonta\|k_Metazoa\|-_Eumetazoa |
| | 1.03578 | 709 | F | 30102 | f_Cicadellidae | -_root\|-_cellular organisms\|d_Eukaryota\|-_Opisthokonta\|k_Metazoa\|-_Eumetazoa |
| | 0.352077 | 241 | F | 139606 | f_Clastopteridae | -_root\|-_cellular organisms\|d_Eukaryota\|-_Opisthokonta\|k_Metazoa\|-_Eumetazoa |
| | 0.0482097 | 33 | F | 33362 | f_Delphacidae | -_root\|-_cellular organisms\|d_Eukaryota\|-_Opisthokonta\|k_Metazoa\|-_Eumetazoa |
| | 0.460183 | 315 | F | 27479 | f_Reduviidae | -_root\|-_cellular organisms\|d_Eukaryota\|-_Opisthokonta\|k_Metazoa\|-_Eumetazoa |
| | 0.306789 | 210 | F | 30083 | f_Miridae | -_root\|-_cellular organisms\|d_Eukaryota\|-_Opisthokonta\|k_Metazoa\|-_Eumetazoa |
| | 0.674935 | 462 | F | 27482 | f_Aphididae | -_root\|-_cellular organisms\|d_Eukaryota\|-_Opisthokonta\|k_Metazoa\|-_Eumetazoa |
| | 0.436809 | 299 | F | 1585420 | f_Liviidae | -_root\|-_cellular organisms\|d_Eukaryota\|-_Opisthokonta\|k_Metazoa\|-_Eumetazoa |
| | 0.495245 | 339 | F | 121221 | f_Pediculidae | -_root\|-_cellular organisms\|d_Eukaryota\|-_Opisthokonta\|k_Metazoa\|-_Eumetazoa |
| | 0.124176 | 85 | F | 48704 | f_Entomobryidae | -_root\|-_cellular organisms\|d_Eukaryota\|-_Opisthokonta\|k_Metazoa\|-_Eumetazoa |
| | 0.0876539 | 60 | F | 36141 | f_Isotomidae | -_root\|-_cellular organisms\|d_Eukaryota\|-_Opisthokonta\|k_Metazoa\|-_Eumetazoa |
| | 0.582899 | 399 | F | 77658 | f_Daphniidae | -_root\|-_cellular organisms\|d_Eukaryota\|-_Opisthokonta\|k_Metazoa\|-_Eumetazoa |
| | 0.270266 | 185 | F | 6757 | f_Portunidae | -_root\|-_cellular organisms\|d_Eukaryota\|-_Opisthokonta\|k_Metazoa\|-_Eumetazoa |
| | 0.0409052 | 28 | F | 134545 | f_Lysianassidae | -_root\|-_cellular organisms\|d_Eukaryota\|-_Opisthokonta\|k_Metazoa\|-_Eumetazoa |
| | 0.159238 | 109 | F | 126955 | f_Linotaeniidae | -_root\|-_cellular organisms\|d_Eukaryota\|-_Opisthokonta\|k_Metazoa\|-_Eumetazoa |
| | 0.863391 | 591 | F | 6939 | f_Ixodidae | -_root\|-_cellular organisms\|d_Eukaryota\|-_Opisthokonta\|k_Metazoa\|-_Eumetazoa |
| | 0.0934975 | 64 | F | 6936 | f_Argasidae | -_root\|-_cellular organisms\|d_Eukaryota\|-_Opisthokonta\|k_Metazoa\|-_Eumetazoa |
| | 0.0350616 | 24 | F | 297309 | f_Nuttalliellidae | -_root\|-_cellular organisms\|d_Eukaryota\|-_Opisthokonta\|k_Metazoa\|-_Eumetazoa |
| | 0.0189917 | 13 | F | 48831 | f_Laelapidae | -_root\|-_cellular organisms\|d_Eukaryota\|-_Opisthokonta\|k_Metazoa\|-_Eumetazoa |
| | 0.0189917 | 13 | F | 6952 | f_Pyroglyphidae | -_root\|-_cellular organisms\|d_Eukaryota\|-_Opisthokonta\|k_Metazoa\|-_Eumetazoa |
| | 0.0219135 | 15 | F | 32262 | f_Tetranychidae | -_root\|-_cellular organisms\|d_Eukaryota\|-_Opisthokonta\|k_Metazoa\|-_Eumetazoa |
| | 0.699771 | 479 | F | 175333 | f_Eresidae | -_root\|-_cellular organisms\|d_Eukaryota\|-_Opisthokonta\|k_Metazoa\|-_Eumetazoa |
| | 0.116872 | 80 | F | 450948 | f_Nephilidae | -_root\|-_cellular organisms\|d_Eukaryota\|-_Opisthokonta\|k_Metazoa\|-_Eumetazoa |
| | 0.043827 | 30 | F | 34643 | f_Theridiidae | -_root\|-_cellular organisms\|d_Eukaryota\|-_Opisthokonta\|k_Metazoa\|-_Eumetazoa |
| | 0.0686623 | 47 | F | 6246 | f_Strongyloididae | -_root\|-_cellular organisms\|d_Eukaryota\|-_Opisthokonta\|k_Metazoa\|-_Eumetazoa |
| | 0.0175308 | 12 | F | 33253 | f_Steinernematida | -_root\|-_cellular organisms\|d_Eukaryota\|-_Opisthokonta\|k_Metazoa\|-_Eumetazoa |
| | 0.0248353 | 17 | F | 6296 | f_Onchocercidae | -_root\|-_cellular organisms\|d_Eukaryota\|-_Opisthokonta\|k_Metazoa\|-_Eumetazoa |
| | 0.0350616 | 24 | F | 6243 | f_Rhabditidae | -_root\|-_cellular organisms\|d_Eukaryota\|-_Opisthokonta\|k_Metazoa\|-_Eumetazoa |
| | 0.124176 | 85 | F | 126387 | f_Haemonchidae | -_root\|-_cellular organisms\|d_Eukaryota\|-_Opisthokonta\|k_Metazoa\|-_Eumetazoa |
| | 0.0262962 | 18 | F | 33278 | f_Ancylostomatida | -_root\|-_cellular organisms\|d_Eukaryota\|-_Opisthokonta\|k_Metazoa\|-_Eumetazoa |
| | 0.0189917 | 13 | F | 2301471 | f_Heterorhabditid | -_root\|-_cellular organisms\|d_Eukaryota\|-_Opisthokonta\|k_Metazoa\|-_Eumetazoa |
| | 0.277571 | 190 | F | 6332 | f_Trichinellidae | -_root\|-_cellular organisms\|d_Eukaryota\|-_Opisthokonta\|k_Metazoa\|-_Eumetazoa |
| | 0.0598969 | 41 | F | 119093 | f_Trichuridae | -_root\|-_cellular organisms\|d_Eukaryota\|-_Opisthokonta\|k_Metazoa\|-_Eumetazoa |
| | 0.0277571 | 19 | F | 6540 | f_Arionidae | -_root\|-_cellular organisms\|d_Eukaryota\|-_Opisthokonta\|k_Metazoa\|-_Eumetazoa |
| | 0.0248353 | 17 | F | 6524 | f_Planorbidae | -_root\|-_cellular organisms\|d_Eukaryota\|-_Opisthokonta\|k_Metazoa\|-_Eumetazoa |
| | 0.0409052 | 28 | F | 54973 | f_Ampullariidae | -_root\|-_cellular organisms\|d_Eukaryota\|-_Opisthokonta\|k_Metazoa\|-_Eumetazoa |
| | 0.0409052 | 28 | F | 69676 | f_Lottiidae | -_root\|-_cellular organisms\|d_Eukaryota\|-_Opisthokonta\|k_Metazoa\|-_Eumetazoa |
| | 0.0306789 | 21 | F | 6563 | f_Ostreidae | -_root\|-_cellular organisms\|d_Eukaryota\|-_Opisthokonta\|k_Metazoa\|-_Eumetazoa |
| | 0.0189917 | 13 | F | 6566 | f_Pectinidae | -_root\|-_cellular organisms\|d_Eukaryota\|-_Opisthokonta\|k_Metazoa\|-_Eumetazoa |
| | 0.0379834 | 26 | F | 6647 | f_Octopodidae | -_root\|-_cellular organisms\|d_Eukaryota\|-_Opisthokonta\|k_Metazoa\|-_Eumetazoa |
| | 0.0175308 | 12 | F | 6214 | f_Hymenolepidida | -_root\|-_cellular organisms\|d_Eukaryota\|-_Opisthokonta\|k_Metazoa\|-_Eumetazoa |
| | 0.0233744 | 16 | F | 31245 | f_Schistosomatida | -_root\|-_cellular organisms\|d_Eukaryota\|-_Opisthokonta\|k_Metazoa\|-_Eumetazoa |
| | 0.0175308 | 12 | F | 6196 | f_Opisthorchiidae | -_root\|-_cellular organisms\|d_Eukaryota\|-_Opisthokonta\|k_Metazoa\|-_Eumetazoa |
| | 0.0482097 | 33 | F | 6407 | f_Glossiphoniidae | -_root\|-_cellular organisms\|d_Eukaryota\|-_Opisthokonta\|k_Metazoa\|-_Eumetazoa |
| | 0.0233744 | 16 | F | 51292 | f_Capitellidae | -_root\|-_cellular organisms\|d_Eukaryota\|-_Opisthokonta\|k_Metazoa\|-_Eumetazoa |
| | 0.0496706 | 34 | F | 33491 | f_Lingulidae | -_root\|-_cellular organisms\|d_Eukaryota\|-_Opisthokonta\|k_Metazoa\|-_Eumetazoa |
| | 0.467488 | 320 | F | 31181 | f_Strongylocentro | -_root\|-_cellular organisms\|d_Eukaryota\|-_Opisthokonta\|k_Metazoa\|-_Eumetazoa |
| | 0.0423661 | 29 | F | 7687 | f_Stichopodidae | -_root\|-_cellular organisms\|d_Eukaryota\|-_Opisthokonta\|k_Metazoa\|-_Eumetazoa |
| | 0.40613 | 278 | F | 35587 | f_Myxobolidae | -_root\|-_cellular organisms\|d_Eukaryota\|-_Opisthokonta\|k_Metazoa\|-_Eumetazoa |
| | 0.0277571 | 19 | F | 46729 | f_Pocilloporidae | -_root\|-_cellular organisms\|d_Eukaryota\|-_Opisthokonta\|k_Metazoa\|-_Eumetazoa |
| | 0.141707 | 97 | F | 1131492 | f_Aspergillaceae | -_root\|-_cellular organisms\|d_Eukaryota\|-_Opisthokonta\|k_Fungi\|-_Dikarya\|p_Asc |
| | 0.0350616 | 24 | F | 27974 | f_Nosematidae | -_root\|-_cellular organisms\|d_Eukaryota\|-_Opisthokonta\|k_Fungi\|-_Fungi incertae |

| Predator | Rel. abundance | CladeReads | TaxRank | TaxID | Name | TaxLineage |
|---|---|---|---|---|---|---|
| **Damselfly** | 66.8842 | 106381 | D | 2759 | d_Eukaryota | -_root\|-_cellular organisms\|d_Eukaryota |
| | 7.8246 | 12757 | F | 27479 | f_Reduviidae | -_root\|-_cellular organisms\|d_Eukaryota\|-_Opisthokonta\|k_Metazoa\|-_Eumetazoa |
| | 2.00016 | 3261 | F | 30083 | f_Miridae | -_root\|-_cellular organisms\|d_Eukaryota\|-_Opisthokonta\|k_Metazoa\|-_Eumetazoa |
| | 1.14637 | 1869 | F | 7533 | f_Lygaeidae | -_root\|-_cellular organisms\|d_Eukaryota\|-_Opisthokonta\|k_Metazoa\|-_Eumetazoa |
| | 0.0601091 | 98 | F | 41702 | f_Alydidae | -_root\|-_cellular organisms\|d_Eukaryota\|-_Opisthokonta\|k_Metazoa\|-_Eumetazoa |
| | 0.0472285 | 77 | F | 50616 | f_Notonectidae | -_root\|-_cellular organisms\|d_Eukaryota\|-_Opisthokonta\|k_Metazoa\|-_Eumetazoa |
| | 1.7217 | 2807 | F | 30102 | f_Cicadellidae | -_root\|-_cellular organisms\|d_Eukaryota\|-_Opisthokonta\|k_Metazoa\|-_Eumetazoa |
| | 0.59557 | 971 | F | 139606 | f_Clastopteridae | -_root\|-_cellular organisms\|d_Eukaryota\|-_Opisthokonta\|k_Metazoa\|-_Eumetazoa |
| | 0.0331213 | 54 | F | 33362 | f_Delphacidae | -_root\|-_cellular organisms\|d_Eukaryota\|-_Opisthokonta\|k_Metazoa\|-_Eumetazoa |
| | 1.36595 | 2227 | F | 27482 | f_Aphididae | -_root\|-_cellular organisms\|d_Eukaryota\|-_Opisthokonta\|k_Metazoa\|-_Eumetazoa |
| | 0.690641 | 1126 | F | 1585420 | f_Liviidae | -_root\|-_cellular organisms\|d_Eukaryota\|-_Opisthokonta\|k_Metazoa\|-_Eumetazoa |
| | 0.0466152 | 76 | F | 121221 | f_Pediculidae | -_root\|-_cellular organisms\|d_Eukaryota\|-_Opisthokonta\|k_Metazoa\|-_Eumetazoa |
| | 1.94618 | 3173 | F | 36668 | f_Formicidae | -_root\|-_cellular organisms\|d_Eukaryota\|-_Opisthokonta\|k_Metazoa\|-_Eumetazoa |
| | 0.223262 | 364 | F | 7458 | f_Apidae | -_root\|-_cellular organisms\|d_Eukaryota\|-_Opisthokonta\|k_Metazoa\|-_Eumetazoa |
| | 0.0441618 | 72 | F | 77572 | f_Halictidae | -_root\|-_cellular organisms\|d_Eukaryota\|-_Opisthokonta\|k_Metazoa\|-_Eumetazoa |
| | 0.625012 | 1019 | F | 7423 | f_Pteromalidae | -_root\|-_cellular organisms\|d_Eukaryota\|-_Opisthokonta\|k_Metazoa\|-_Eumetazoa |
| | 0.0662426 | 108 | F | 7402 | f_Braconidae | -_root\|-_cellular organisms\|d_Eukaryota\|-_Opisthokonta\|k_Metazoa\|-_Eumetazoa |
| | 0.835393 | 1362 | F | 7392 | f_Glossinidae | -_root\|-_cellular organisms\|d_Eukaryota\|-_Opisthokonta\|k_Metazoa\|-_Eumetazoa |
| | 0.0337347 | 55 | F | 7371 | f_Calliphoridae | -_root\|-_cellular organisms\|d_Eukaryota\|-_Opisthokonta\|k_Metazoa\|-_Eumetazoa |
| | 0.0153339 | 25 | F | 7366 | f_Muscidae | -_root\|-_cellular organisms\|d_Eukaryota\|-_Opisthokonta\|k_Metazoa\|-_Eumetazoa |
| | 0.379668 | 619 | F | 7211 | f_Tephritidae | -_root\|-_cellular organisms\|d_Eukaryota\|-_Opisthokonta\|k_Metazoa\|-_Eumetazoa |
| | 0.405429 | 661 | F | 7205 | f_Tabanidae | -_root\|-_cellular organisms\|d_Eukaryota\|-_Opisthokonta\|k_Metazoa\|-_Eumetazoa |
| | 0.314653 | 513 | F | 7157 | f_Culicidae | -_root\|-_cellular organisms\|d_Eukaryota\|-_Opisthokonta\|k_Metazoa\|-_Eumetazoa |
| | 0.00613358 | 10 | F | 41811 | f_Chaoboridae | -_root\|-_cellular organisms\|d_Eukaryota\|-_Opisthokonta\|k_Metazoa\|-_Eumetazoa |
| | 0.0042935 | 7 | F | 7149 | f_Chironomidae | -_root\|-_cellular organisms\|d_Eukaryota\|-_Opisthokonta\|k_Metazoa\|-_Eumetazoa |
| | 0.0453885 | 74 | F | 7197 | f_Psychodidae | -_root\|-_cellular organisms\|d_Eukaryota\|-_Opisthokonta\|k_Metazoa\|-_Eumetazoa |
| | 0.583303 | 951 | F | 7049 | f_Lampyridae | -_root\|-_cellular organisms\|d_Eukaryota\|-_Opisthokonta\|k_Metazoa\|-_Eumetazoa |
| | 0.0502953 | 82 | F | 50527 | f_Buprestidae | -_root\|-_cellular organisms\|d_Eukaryota\|-_Opisthokonta\|k_Metazoa\|-_Eumetazoa |
| | 0.135552 | 221 | F | 7065 | f_Tenebrionidae | -_root\|-_cellular organisms\|d_Eukaryota\|-_Opisthokonta\|k_Metazoa\|-_Eumetazoa |
| | 0.134939 | 220 | F | 34667 | f_Cerambycidae | -_root\|-_cellular organisms\|d_Eukaryota\|-_Opisthokonta\|k_Metazoa\|-_Eumetazoa |
| | 0.0441618 | 72 | F | 7042 | f_Curculionidae | -_root\|-_cellular organisms\|d_Eukaryota\|-_Opisthokonta\|k_Metazoa\|-_Eumetazoa |
| | 0.0122672 | 20 | F | 7055 | f_Scarabaeidae | -_root\|-_cellular organisms\|d_Eukaryota\|-_Opisthokonta\|k_Metazoa\|-_Eumetazoa |
| | 0.386415 | 630 | F | 7100 | f_Noctuidae | -_root\|-_cellular organisms\|d_Eukaryota\|-_Opisthokonta\|k_Metazoa\|-_Eumetazoa |
| | 0.066856 | 109 | F | 33415 | f_Nymphalidae | -_root\|-_cellular organisms\|d_Eukaryota\|-_Opisthokonta\|k_Metazoa\|-_Eumetazoa |
| | 0.0263744 | 43 | F | 7143 | f_Papilionidae | -_root\|-_cellular organisms\|d_Eukaryota\|-_Opisthokonta\|k_Metazoa\|-_Eumetazoa |
| | 0.075443 | 123 | F | 7089 | f_Bombycidae | -_root\|-_cellular organisms\|d_Eukaryota\|-_Opisthokonta\|k_Metazoa\|-_Eumetazoa |
| | 0.0184007 | 30 | F | 82593 | f_Geometridae | -_root\|-_cellular organisms\|d_Eukaryota\|-_Opisthokonta\|k_Metazoa\|-_Eumetazoa |
| | 0.0147206 | 24 | F | 32434 | f_Gelechiidae | -_root\|-_cellular organisms\|d_Eukaryota\|-_Opisthokonta\|k_Metazoa\|-_Eumetazoa |
| | 0.76179 | 1242 | F | 46562 | f_Kalotermitidae | -_root\|-_cellular organisms\|d_Eukaryota\|-_Opisthokonta\|k_Metazoa\|-_Eumetazoa |
| | 0.123898 | 202 | F | 7501 | f_Termopsidae | -_root\|-_cellular organisms\|d_Eukaryota\|-_Opisthokonta\|k_Metazoa\|-_Eumetazoa |
| | 0.51154 | 834 | F | 1049651 | f_Ectobiidae | -_root\|-_cellular organisms\|d_Eukaryota\|-_Opisthokonta\|k_Metazoa\|-_Eumetazoa |
| | 0.0564289 | 92 | F | 36141 | f_Isotomidae | -_root\|-_cellular organisms\|d_Eukaryota\|-_Opisthokonta\|k_Metazoa\|-_Eumetazoa |
| | 0.00797365 | 13 | F | 48704 | f_Entomobryidae | -_root\|-_cellular organisms\|d_Eukaryota\|-_Opisthokonta\|k_Metazoa\|-_Eumetazoa |
| | 0.0594957 | 97 | F | 72034 | f_Caligidae | -_root\|-_cellular organisms\|d_Eukaryota\|-_Opisthokonta\|k_Metazoa\|-_Eumetazoa |
| | 0.0355747 | 58 | F | 134545 | f_Lysianassidae | -_root\|-_cellular organisms\|d_Eukaryota\|-_Opisthokonta\|k_Metazoa\|-_Eumetazoa |
| | 0.0110404 | 18 | F | 6757 | f_Portunidae | -_root\|-_cellular organisms\|d_Eukaryota\|-_Opisthokonta\|k_Metazoa\|-_Eumetazoa |
| | 0.0662426 | 108 | F | 77658 | f_Daphniidae | -_root\|-_cellular organisms\|d_Eukaryota\|-_Opisthokonta\|k_Metazoa\|-_Eumetazoa |
| | 0.0116538 | 19 | F | 126955 | f_Linotaeniidae | -_root\|-_cellular organisms\|d_Eukaryota\|-_Opisthokonta\|k_Metazoa\|-_Eumetazoa |
| | 0.534235 | 871 | F | 450948 | f_Nephilidae | -_root\|-_cellular organisms\|d_Eukaryota\|-_Opisthokonta\|k_Metazoa\|-_Eumetazoa |
| | 0.034348 | 56 | F | 34643 | f_Theridiidae | -_root\|-_cellular organisms\|d_Eukaryota\|-_Opisthokonta\|k_Metazoa\|-_Eumetazoa |
| | 0.273558 | 446 | F | 6936 | f_Argasidae | -_root\|-_cellular organisms\|d_Eukaryota\|-_Opisthokonta\|k_Metazoa\|-_Eumetazoa |
| | 0.118991 | 194 | F | 6939 | f_Ixodidae | -_root\|-_cellular organisms\|d_Eukaryota\|-_Opisthokonta\|k_Metazoa\|-_Eumetazoa |
| | 0.0778964 | 127 | F | 297309 | f_Nuttalliellidae | -_root\|-_cellular organisms\|d_Eukaryota\|-_Opisthokonta\|k_Metazoa\|-_Eumetazoa |
| | 0.0294412 | 48 | F | 6952 | f_Pyroglyphidae | -_root\|-_cellular organisms\|d_Eukaryota\|-_Opisthokonta\|k_Metazoa\|-_Eumetazoa |
| | 0.228169 | 372 | F | 6246 | f_Strongyloididae | -_root\|-_cellular organisms\|d_Eukaryota\|-_Opisthokonta\|k_Metazoa\|-_Eumetazoa |
| | 0.0251477 | 41 | F | 33253 | f_Steinernematida | -_root\|-_cellular organisms\|d_Eukaryota\|-_Opisthokonta\|k_Metazoa\|-_Eumetazoa |
| | 0.0521354 | 85 | F | 33823 | f_Aphelenchoidid | -_root\|-_cellular organisms\|d_Eukaryota\|-_Opisthokonta\|k_Metazoa\|-_Eumetazoa |
| | 0.0533621 | 87 | F | 6243 | f_Rhabditidae | -_root\|-_cellular organisms\|d_Eukaryota\|-_Opisthokonta\|k_Metazoa\|-_Eumetazoa |
| | 0.0116538 | 19 | F | 6296 | f_Onchocercidae | -_root\|-_cellular organisms\|d_Eukaryota\|-_Opisthokonta\|k_Metazoa\|-_Eumetazoa |
| | 0.0852567 | 139 | F | 126387 | f_Haemonchidae | -_root\|-_cellular organisms\|d_Eukaryota\|-_Opisthokonta\|k_Metazoa\|-_Eumetazoa |
| | 0.00858701 | 14 | F | 2301471 | f_Heterorhabditid | -_root\|-_cellular organisms\|d_Eukaryota\|-_Opisthokonta\|k_Metazoa\|-_Eumetazoa |
| | 0.106111 | 173 | F | 119093 | f_Trichuridae | -_root\|-_cellular organisms\|d_Eukaryota\|-_Opisthokonta\|k_Metazoa\|-_Eumetazoa |
| | 0.0398683 | 65 | F | 6332 | f_Trichinellidae | -_root\|-_cellular organisms\|d_Eukaryota\|-_Opisthokonta\|k_Metazoa\|-_Eumetazoa |
| | 0.16254 | 265 | F | 6524 | f_Planorbidae | -_root\|-_cellular organisms\|d_Eukaryota\|-_Opisthokonta\|k_Metazoa\|-_Eumetazoa |
| | 0.0300545 | 49 | F | 6540 | f_Arionidae | -_root\|-_cellular organisms\|d_Eukaryota\|-_Opisthokonta\|k_Metazoa\|-_Eumetazoa |
| | 0.0104271 | 17 | F | 54973 | f_Ampullariidae | -_root\|-_cellular organisms\|d_Eukaryota\|-_Opisthokonta\|k_Metazoa\|-_Eumetazoa |
| | 0.00674693 | 11 | F | 6489 | f_Conidae | -_root\|-_cellular organisms\|d_Eukaryota\|-_Opisthokonta\|k_Metazoa\|-_Eumetazoa |
| | 0.0147206 | 24 | F | 69676 | f_Lottiidae | -_root\|-_cellular organisms\|d_Eukaryota\|-_Opisthokonta\|k_Metazoa\|-_Eumetazoa |
| | 0.150273 | 245 | F | 6647 | f_Octopodidae | -_root\|-_cellular organisms\|d_Eukaryota\|-_Opisthokonta\|k_Metazoa\|-_Eumetazoa |
| | 0.0159473 | 26 | F | 6196 | f_Opisthorchiidae | -_root\|-_cellular organisms\|d_Eukaryota\|-_Opisthokonta\|k_Metazoa\|-_Eumetazoa |
| | 0.0147206 | 24 | F | 31245 | f_Schistosomatida | -_root\|-_cellular organisms\|d_Eukaryota\|-_Opisthokonta\|k_Metazoa\|-_Eumetazoa |
| | 0.0392549 | 64 | F | 39216 | f_Macrostomidae | -_root\|-_cellular organisms\|d_Eukaryota\|-_Opisthokonta\|k_Metazoa\|-_Eumetazoa |
| | 0.0134939 | 22 | F | 6214 | f_Hymenolepidida | -_root\|-_cellular organisms\|d_Eukaryota\|-_Opisthokonta\|k_Metazoa\|-_Eumetazoa |
| | 0.0736029 | 120 | F | 6407 | f_Glossiphoniidae | -_root\|-_cellular organisms\|d_Eukaryota\|-_Opisthokonta\|k_Metazoa\|-_Eumetazoa |
| | 0.00674693 | 11 | F | 51292 | f_Capitellidae | -_root\|-_cellular organisms\|d_Eukaryota\|-_Opisthokonta\|k_Metazoa\|-_Eumetazoa |
| | 0.0196274 | 32 | F | 31181 | f_Strongylocentro | -_root\|-_cellular organisms\|d_Eukaryota\|-_Opisthokonta\|k_Metazoa\|-_Eumetazoa |
| | 0.0110404 | 18 | F | 7687 | f_Stichopodidae | -_root\|-_cellular organisms\|d_Eukaryota\|-_Opisthokonta\|k_Metazoa\|-_Eumetazoa |
| | 0.0760564 | 124 | F | 6080 | f_Hydridae | -_root\|-_cellular organisms\|d_Eukaryota\|-_Opisthokonta\|k_Metazoa\|-_Eumetazoa |
| | 0.0220809 | 36 | F | 35587 | f_Myxobolidae | -_root\|-_cellular organisms\|d_Eukaryota\|-_Opisthokonta\|k_Metazoa\|-_Eumetazoa |
| | 0.0110404 | 18 | F | 46729 | f_Pocilloporidae | -_root\|-_cellular organisms\|d_Eukaryota\|-_Opisthokonta\|k_Metazoa\|-_Eumetazoa |
| | 0.00858701 | 14 | F | 1506295 | f_Serendipitaceae | -_root\|-_cellular organisms\|d_Eukaryota\|-_Opisthokonta\|k_Fungi\|-_Dikarya\|p_Bas |
| | 0.00981372 | 16 | F | 1799696 | f_Sporidiobolacea | -_root\|-_cellular organisms\|d_Eukaryota\|-_Opisthokonta\|k_Fungi\|-_Dikarya\|p_Bas |
| | 0.0153339 | 25 | F | 34371 | f_Erysiphaceae | -_root\|-_cellular organisms\|d_Eukaryota\|-_Opisthokonta\|k_Fungi\|-_Dikarya\|p_Asc |
| | 0.00981372 | 16 | F | 1131492 | f_Aspergillaceae | -_root\|-_cellular organisms\|d_Eukaryota\|-_Opisthokonta\|k_Fungi\|-_Dikarya\|p_Asc |
| | 0.017174 | 28 | F | 27974 | f_Nosematidae | -_root\|-_cellular organisms\|d_Eukaryota\|-_Opisthokonta\|k_Fungi\|-_Fungi incertae |
| | 0.00674693 | 11 | F | 35232 | f_Pleistophoridae | -_root\|-_cellular organisms\|d_Eukaryota\|-_Opisthokonta\|k_Fungi\|-_Fungi incertae |

| Predator | Rel. abundance | CladeReads | TaxRank | TaxID | Name | TaxLineage |
|---|---|---|---|---|---|---|
| **Mosquito fish (*G.*** | 94.0526 | 7834 | D | 2759 | d_Eukaryota | -_root\|-_cellular organisms\|d_Eukaryota |
| | 0.0204498 | 35 | F | 31181 | f_Strongylocentro | -_root\|-_cellular organisms\|d_Eukaryota\|-_Opisthokonta\|k_Metazoa\|-_Eumetazoa |
| | 0.014607 | 25 | F | 7687 | f_Stichopodidae | -_root\|-_cellular organisms\|d_Eukaryota\|-_Opisthokonta\|k_Metazoa\|-_Eumetazoa |
| | 0.583111 | 998 | F | 6208 | f_Taeniidae | -_root\|-_cellular organisms\|d_Eukaryota\|-_Opisthokonta\|k_Metazoa\|-_Eumetazoa |
| | 0.00642707 | 11 | F | 28843 | f_Diphyllobothriid | -_root\|-_cellular organisms\|d_Eukaryota\|-_Opisthokonta\|k_Metazoa\|-_Eumetazoa |
| | 0.014607 | 25 | F | 31245 | f_Schistosomatida | -_root\|-_cellular organisms\|d_Eukaryota\|-_Opisthokonta\|k_Metazoa\|-_Eumetazoa |
| | 0.0461581 | 79 | F | 54973 | f_Ampullariidae | -_root\|-_cellular organisms\|d_Eukaryota\|-_Opisthokonta\|k_Metazoa\|-_Eumetazoa |
| | 0.00876419 | 15 | F | 33491 | f_Lingulidae | -_root\|-_cellular organisms\|d_Eukaryota\|-_Opisthokonta\|k_Metazoa\|-_Eumetazoa |
| | 0.00584279 | 10 | F | 51292 | f_Capitellidae | -_root\|-_cellular organisms\|d_Eukaryota\|-_Opisthokonta\|k_Metazoa\|-_Eumetazoa |
| | 0.00876419 | 15 | F | 7392 | f_Glossinidae | -_root\|-_cellular organisms\|d_Eukaryota\|-_Opisthokonta\|k_Metazoa\|-_Eumetazoa |
| | 0.00584279 | 10 | F | 7371 | f_Calliphoridae | -_root\|-_cellular organisms\|d_Eukaryota\|-_Opisthokonta\|k_Metazoa\|-_Eumetazoa |
| | 0.0128541 | 22 | F | 7157 | f_Culicidae | -_root\|-_cellular organisms\|d_Eukaryota\|-_Opisthokonta\|k_Metazoa\|-_Eumetazoa |
| | 0.00701135 | 12 | F | 36668 | f_Formicidae | -_root\|-_cellular organisms\|d_Eukaryota\|-_Opisthokonta\|k_Metazoa\|-_Eumetazoa |
| | 0.00817991 | 14 | F | 27482 | f_Aphididae | -_root\|-_cellular organisms\|d_Eukaryota\|-_Opisthokonta\|k_Metazoa\|-_Eumetazoa |
| | 0.00759563 | 13 | F | 30102 | f_Cicadellidae | -_root\|-_cellular organisms\|d_Eukaryota\|-_Opisthokonta\|k_Metazoa\|-_Eumetazoa |
| | 0.0192812 | 33 | F | 121221 | f_Pediculidae | -_root\|-_cellular organisms\|d_Eukaryota\|-_Opisthokonta\|k_Metazoa\|-_Eumetazoa |
| | 0.0257083 | 44 | F | 77658 | f_Daphniidae | -_root\|-_cellular organisms\|d_Eukaryota\|-_Opisthokonta\|k_Metazoa\|-_Eumetazoa |
| | 0.00584279 | 10 | F | 126955 | f_Linotaeniidae | -_root\|-_cellular organisms\|d_Eukaryota\|-_Opisthokonta\|k_Metazoa\|-_Eumetazoa |
| | 0.00701135 | 12 | F | 6939 | f_Ixodidae | -_root\|-_cellular organisms\|d_Eukaryota\|-_Opisthokonta\|k_Metazoa\|-_Eumetazoa |
| | 0.00759563 | 13 | F | 175333 | f_Eresidae | -_root\|-_cellular organisms\|d_Eukaryota\|-_Opisthokonta\|k_Metazoa\|-_Eumetazoa |
| | 0.0262926 | 45 | F | 318477 | f_Dracunculidae | -_root\|-_cellular organisms\|d_Eukaryota\|-_Opisthokonta\|k_Metazoa\|-_Eumetazoa |
| | 0.00701135 | 12 | F | 6246 | f_Strongyloididae | -_root\|-_cellular organisms\|d_Eukaryota\|-_Opisthokonta\|k_Metazoa\|-_Eumetazoa |
| | 0.00701135 | 12 | F | 126387 | f_Haemonchidae | -_root\|-_cellular organisms\|d_Eukaryota\|-_Opisthokonta\|k_Metazoa\|-_Eumetazoa |
| | 0.0490795 | 84 | F | 6332 | f_Trichinellidae | -_root\|-_cellular organisms\|d_Eukaryota\|-_Opisthokonta\|k_Metazoa\|-_Eumetazoa |
| | 0.0163598 | 28 | F | 119093 | f_Trichuridae | -_root\|-_cellular organisms\|d_Eukaryota\|-_Opisthokonta\|k_Metazoa\|-_Eumetazoa |
| | 0.0157755 | 27 | F | 45349 | f_Edwardsiidae | -_root\|-_cellular organisms\|d_Eukaryota\|-_Opisthokonta\|k_Metazoa\|-_Eumetazoa |
| | 0.0157755 | 27 | F | 46729 | f_Pocilloporidae | -_root\|-_cellular organisms\|d_Eukaryota\|-_Opisthokonta\|k_Metazoa\|-_Eumetazoa |
| | 0.00993275 | 17 | F | 6080 | f_Hydridae | -_root\|-_cellular organisms\|d_Eukaryota\|-_Opisthokonta\|k_Metazoa\|-_Eumetazoa |
